# Rapid Knowledgebase Construction and Hypotheses Generation Using Extractive Literature Search

**DOI:** 10.1101/2022.02.13.480241

**Authors:** Shaked Launer-Wachs, Hillel Taub-Tabib, Yoav Goldberg, Yosi Shamay

## Abstract

As knowledgebases become increasingly important for structuring vast amounts of scientific knowledge and making it accessible to researchers, their construction entails expensive multi-year projects involving teams of bio-curators, computer scientists, or both. This restricts the coverage of existing knowledgebases to a limited set of popular topics, leaving a long tail of more specialized interests uncovered.

We present a methodology and a supporting tool to allow individual researchers or small teams, without background in bio-curation or computer science, to mine the scientific literature and construct ad-hoc, personalized, and literature-anchored knowledgebases, that are tailored around their specific research interests and support their scientific goals. The time investment involved in creating a knowledgebase ranges from a few hours to a few weeks, depending on the desired coverage and accuracy.

We demonstrate the methodology by constructing knowledgebases for different purposes: a high-level overview of challenges and controversies in a field (the cancer frontiers knowledgebase); a mapping of main concepts and interactions in a field, to support lab-internal hypothesis generation (tissue engineering and regeneration, cancer surgery and radiotherapy knowledgebases); and a comprehensive and accurate knowledgebase designated as an online up-to-date resource for the wider research community (the cell specific drug delivery knowledgebase). In each case we show how the structured knowledgebase, coupled with effective visualizations, facilitates effective data exploration, hypothesis generation and meta-analysis.

We implement the method as part of an open source web-based platform for knowledgebase construction, available publicly and freely at https://spike-kbc.apps.allenai.org.

## Main

The overwhelming extent of biomedical data, based on unstructured, narrative papers, books and medical records has made it unfeasible for scientists to keep up with the breadth of published work in their fields of expertise, or enter new fields^1^. Expectedly, a variety of solutions have been developed in recent years to address this challenge, aimed at presenting up-to-date, concise, and yet comprehensive information: Nowadays, databases and ontologies (commonly referred here as knowledgebases) gain popularity as alternative platforms for knowledge organization and presentation, as they provide highly accessible and readily updated representations of contemporary data to let end users interact with entries of interest and their relations, while avoiding irrelevant information^2–4^.

The majority of biomedical knowledgebases are constructed by dedicated teams of bio-curators. These are typically trained Ph.D. level trained individuals who manually review multiple information sources, contact authors where needed, and upon which produce structured ontologies which represent our ‘best view’ of the data. They also ensure that identifiers and links to other ontologies are used where possible and optimize data representation and interoperability^5, 6^. Hence, while biocuration has produced highly utilized resources, the meticulous processes and quality controls involved may preclude many research labs from producing similar knowledgebases ^7, 8^.

As an alternative to biocuration, trained Natural Language Processing (NLP) models can be used for automated knowledgebase construction (AKBC)^9–11^. A prominent example is CancerMine, a cancer-focused knowledgebase which was mined from the scientific literature and is being kept up to date regularly, with minimal human involvement^12^. However, despite its benefits, AKBC involves significant efforts of data collection, model training and tuning. As both biocuration and AKBC demand inter-disciplinary collaborations, their practice is typically out of reach for individual researchers or small labs. Therefore, existing knowledgebases tend to focus on relation types that are expected to be of interest to a wide scientific community, thus leaving a wide “tail” of relations which are not covered by any knowledgebase.

We advocate to supplement such large-scale knowledgebase construction efforts with ad-hoc, personalized, literature-based and literature-anchored knowledgebases that can be rapidly created by individuals and small teams based on their current research interests, and without requiring biocuration or computer science expertise.

Such personalized knowledgebase are meant to be created rapidly, used to explore the literature, discover connections, and form hypotheses. Moreover, as the researcher who created the knowledgebase is also its initial consumer, it can be kept up to date as long as the research is active, and its quality can improve along its lifetime, depending on the researcher’s goals and expectations. A personalized knowledgebase can start as a private, crude effort, providing a high-level view of a field the researcher is just starting to explore, and gradually change into a public, highly precise resource, tracking and expanding the knowledge in a field the researcher is already an expert on.

We show how such ad-hoc knowledgebase can be built by leveraging extractive-search. Extractive-search is an emerging query-based interface to the scientific literature that aims to provide scientists with access to text-mining and NLP building-blocks, without requiring extensive training in NLP, text-mining or computer science, and without requiring programming^13^. In this work, we extend extractive-search with knowledgebase curation tools, resulting in a web-based platform for biomedical knowledgebase construction. The system, named SPIKE-KBC, allows researchers to construct and query personalized knowledgebases on an ad-hoc basis, according to their current research interests.

We demonstrate this approach with three levels of knowledgebase construction: we commence with a general application of extractive search to form a small scale, high-level overview, focusing on abstract challenges in the field of cancer therapy. We then advance to rapid knowledgebase construction when entering a new field, enabling hypothesis generation and exploration (this is demonstrated with knowledgebases on tissue engineering, radiotherapy and surgery). We conclude with a fully annotated, comprehensive public knowledgebase in targeted drug delivery; the authors’ domain of expertise. For each of these workflows, we demonstrate the creation process, how the resulting knowledgebase can be explored, and how scientific insights can be gained.

## Results

### Method and system for the construction of personalized knowledgebases

In this work, we advocate the rapid construction and exploration of *ad-hoc,* literature-based and personalized knowledgebases as part of the scientific workflow, and demonstrate (a) how researchers without experience in NLP can construct such knowledgebases using extractive-search; and (b) how such constructed knowledgebases can be used to gain scientific insights.

The proposed method is based on Extractive Search and on the use of ‘‘extractive queries’’^13^. An extractive query is a query which combines the standard keyword and search operators that are used to select documents, with capture operators that allow to extract values from the selected documents. As a concrete example, given a list of Chemicals and a list of Drugs, one can form a query that counts all the joint occurrences of a chemical and a drug in a sentence that is mentioned in a paragraph that contains the term *treatment*. Beyond term co-occurrence, extractive queries can impose additional linear, syntactic or semantic restrictions on matching texts^14, 15^.

At a high level, the method consists of the following steps (**Fig 1,** graphic workflow): A user decides on the set of relations they want to capture; the user collects lists of entities of interest, using a combination of pre-defined ontologies (if these exist) and extractive queries over the literature; the user formulates a list of extractive queries that suggest a relation between the entities; optionally, the user verifies the resulting extractions; at this point we have a mostly-correct knowledgebase that is linked to the literature; the user then queries/browses the knowledgebase to explore the literature and form hypotheses. While doing so, the user can revise the extractive queries and entity lists to refine or expand the knowledgebase. We’ll now discuss these steps in more detail.

**Figure 1.**
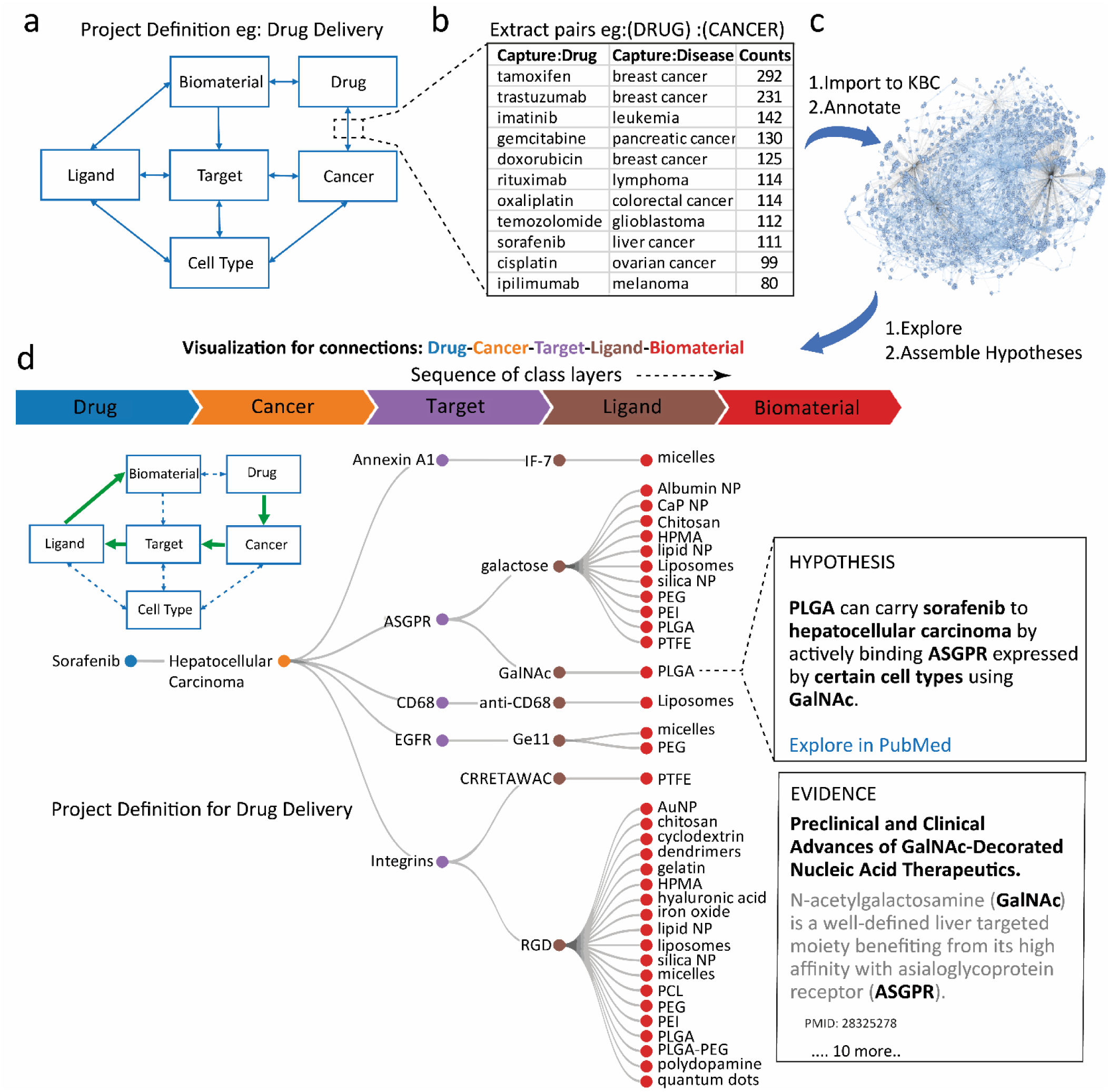
knowledgebase construction using extractive search. **a,** Example of project definitions based on entity classes and defined relation pairs. **b,** Example of pairs extraction using SPIKE for Cancer-Drug relation. **c,** Final steps of knowledgebase construction are importing pairs extraction queries and annotating pair relation instances. **d,** knowledgebase user interface visualization for hypothesis exploration with 5 class layers filtered for sorafenib (drug) and hepatocellular carcinoma (cancer).

#### 1. Deciding on a set of relations

This step is derived from the scientific question at hand, and the intended use of the knowledge-base. It can span a single relation (‘‘which ligands are associated with which biomaterial’’) but richer knowledge-bases can be formed by considering a network of linked relations, as in **Fig 1a**.

#### 2. Collecting Name Lists for Entities

The vast majority of relations hold between typed entities (e.g. ‘‘ligands’’ and ‘‘biomaterias’’), and the initial stage requires identifying literature occurrences of these entities. Name lists representing the different entity types can often be downloaded from online sources or obtained from existing lab records. When no relevant source is available, a standard set of extractive queries can be used to bootstrap an effective list. **Table 1** shows a set of queries we used to obtain biomedical polymers, these queries, based on generic Hearst Patterns^16–18^, can be trivially modified to extract other types of entities directly from the scientific literature.

#### 3. Extracting candidate relation instances

Once entity lists are compiled, extractive queries are used to find potential pairwise connections between entities (relation instances). Researches can use basic queries, e.g. queries to retrieve pairs of ligands and targets which appear in the same sentence, or more involved ones, e.g. queries which retrieve pairs of drugs and cancers which are in close structural proximity to one another (**Fig 1b**). With deeper knowledge of extractive search, more sophisticated queries can be formed, and precision can be increased. However, we find that co-occurrence queries based on domain knowledge are often sufficient to support the annotation step that follows, and that domain experts find these queries intuitive and simple to use.

#### 4. Annotating candidate instances

After importing the queries, researchers are presented with a tabular results view. Each result row in this view consists of a pair of entities and an evidence sentence (i.e. the sentence which matched an extractive query and the captured entities within it). The results are grouped by entity pairs such that results which are associated with the same entity pair appear in sequence. Researchers can label each result row as “positive” or “negative” (whether the evidence indicates that the entity pair satisfies the relation or not). Once a result is labeled, the associated entity pair, evidence sentence and label are added to the knowledgebase. By default, once an entity pair is added, all results associated with it are removed from the results view. This allows researchers to focus on undiscovered connections while skipping established ones. Furthermore, the results are sorted such that entity pairs with more evidence sentences appear before those with less evidence. This allows researchers to prioritize adding frequently reported associations before less established ones.

The annotation and extraction stages are iterative: after labeling some of the results, researchers can choose to refine and re-run the underlying queries. This allows them to improve the recall and precision of unlabeled results without affecting the labeled results already added to the knowledgebase (**Fig 1c**).

#### 5. Visually exploring the knowledgebase and generating novel research hypotheses

The result of the process is a dense graph of relations. Rather than seeing the entire graph at once, we suggest deriving *linear paths of interest* between entities, and visualizing them as a tree. The first layer of the tree is populated with entities of a selected class (e.g., entities of class *Biomaterial*) and subsequent levels are populated by layering more classes, based on the defined relations. Consequently, each layer is linked to the former by relation instances Per instance, if relations are defined between the classes *Biomaterial* and *Ligand* and between the classes *Biomaterial* and *Target,* then entities of either class Ligand or Target can be selected to populate the second layer of the tree, etc. (**Fig 1d**).

This form of visualization generates a sequence of relations between layered classes to form multi-component hypotheses, while each linear, horizontal path along the knowledge tree can be seen as a research hypothesis: pairs of adjacent entities on the path are known to be related to one another, so a hypothesis can be made that all entities on the path might be transitively dependent. As the total number of hypotheses in a given knowledgebase can be very large (>100,000), the user can filter in or out entities from each layer to reduce the hypotheses’ branching. In addition, to allow integration with existing platforms, the results can be exported in Web Ontology Language (OWL format) which can be easily incorporated with other external platforms such as BioPortal and Protégé^19, 20^ or as CSV, facilitating further analysis of the knowledgebase by quantifying entities and frequencies of relations.

### Use case 1: Challenges in Cancer Research

We sought to demonstrate a challenging literature review task of capturing unknown entities and concepts from intangible classes. We thus constructed a knowledgebase to *map challenges in cancer research* using three conceptual classes: ‘controversial’, ‘still unknown/unclear’ and ‘unmet need’. We used structured (syntactic) queries in SPIKE, the extractive search engine used by SPIKE-KBC, to capture entities from structured context. For example, to retrieve controversial findings we used the query “<>:something $remains/still/is $controversial”, restricted to papers that have words pertaining to different cancer types in their abstract and filtered to the last three years to reflect only contemporary topics. The queries yielded entities from diverse biomedical classes from drugs and genes to abstract concepts such as ‘the role of surgery’ or ‘alcohol consumption’ **(Fig 2a)**. Then we queried for these terms together with a list of cancer types to capture matching pairs of cancer type-controversial entities. The results were imported to the SPIKE-KBC annotation module and went through manual triage, rejecting only out-of-context entities, establishing an overview of current challenges mapped by cancer type. **Fig 2b** shows the top ranked cancers and entities by their number of relations which aggregate several insights in cancer research. For example, the most common ‘controversial’ entities for all cancers, are ‘optimal treatment’ and ‘the role of surgery’. Of the different cancer types, the highest number of ‘controversial’ relations applied to breast cancer and hepatocellular carcinoma. We show an example for these types of relations in pancreatic cancer (**Fig 2c**). In the class of ‘still unknown’, the most common entities in multiple cancers were long noncoding RNA (lncRNA), circRNAs and PDL1. We curated 29 entities that are unmet needs, such as ‘novel therapy’ and ‘biomarkers’, of which, ‘breast cancer’ notably had the highest number (17 vs. median of 2). An example for these relations is shown for ovarian cancer in **Fig 2c**. The estimated time for the construction and quantitative evaluation of the knowledgebase was less than 3 hours.

**Figure 2.**
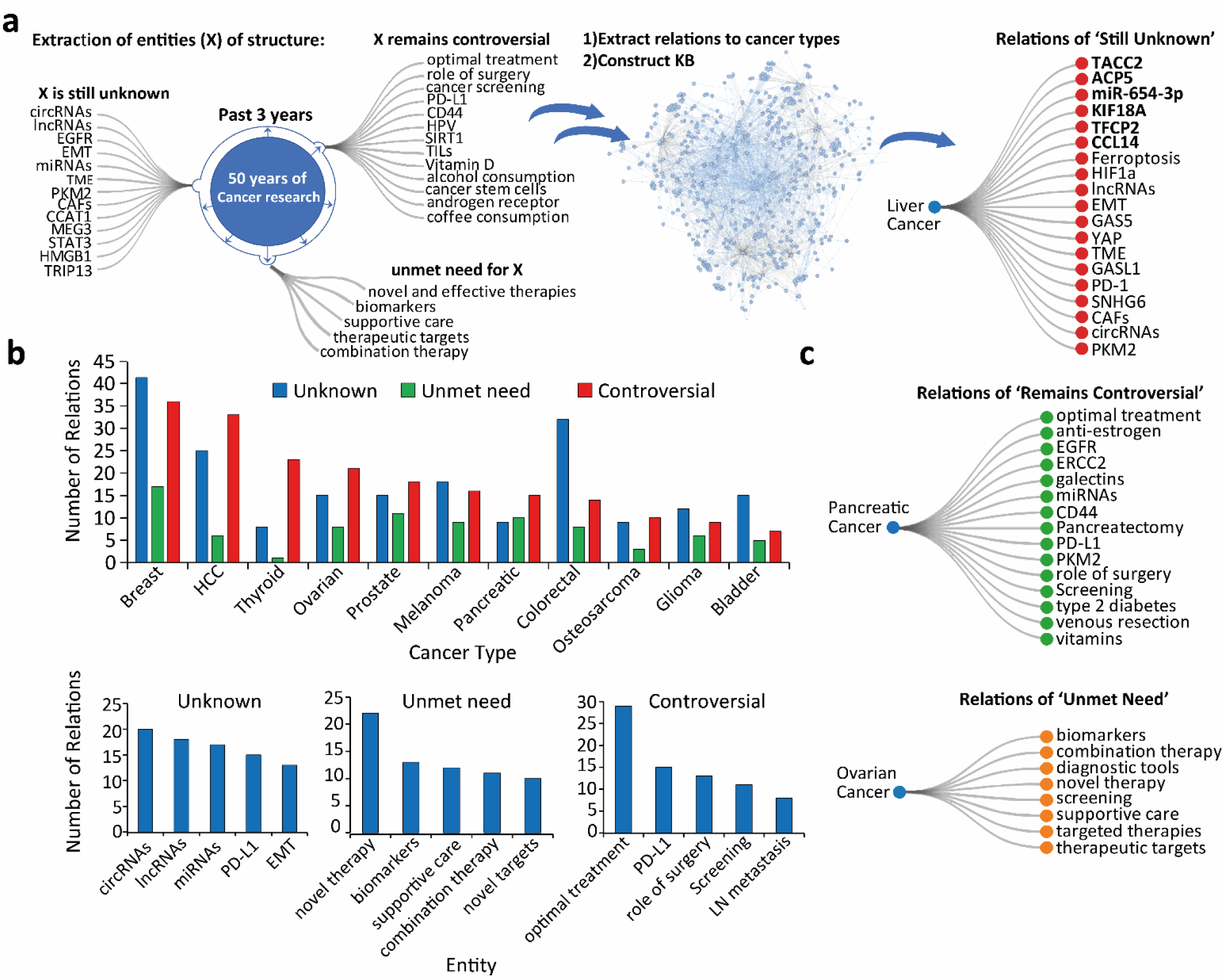
Knowledge Base Construction for Literature Exploration. **a,** knowledgebase construction scheme using three different structural queries for capturing entities in conceptual classes (controversial, unknown, in need). Example of entities identified as unknown or unclear, related to liver cancer with exclusive entities in bold (right). **b,** Top entities in the knowledgebase sorted by number of relations in different cancer types (top) and common to most cancers (bottom). **c**, examples of entities found controversial in relation to pancreatic cancer (top) and example of ‘unmet need’ relations in ovarian cancer.

### Use case 2a: Rapid Hypothesis Generation and Exploration: Tissue Engineering and Regeneration

The field of tissue engineering and regeneration is one of the major disciplines in biomedical engineering and could be a great example for rapid knowledgebase construction. First, we set to define the main hypothesis in this field with a combinatorial relation of classes. Tissue engineering is defined as the use of various cells, material engineering methods and biochemical factors to restore, maintain, improve, or replace biological tissues^21^. We took this simple definition and reconstructed it to the following combinatorial hypothesis of entity classes: A certain medical condition **A** can be treated with a scaffold made from material **B** which can support the growth of cell type **C** by combining bioactive element **D** (growth factors, hormones). We also noted that there many different methods to combine the cells with scaffolds and how to introduce the scaffold to the damaged tissue and for that purpose we added another class Method **E (Fig 3a)**.

**Figure 3.**
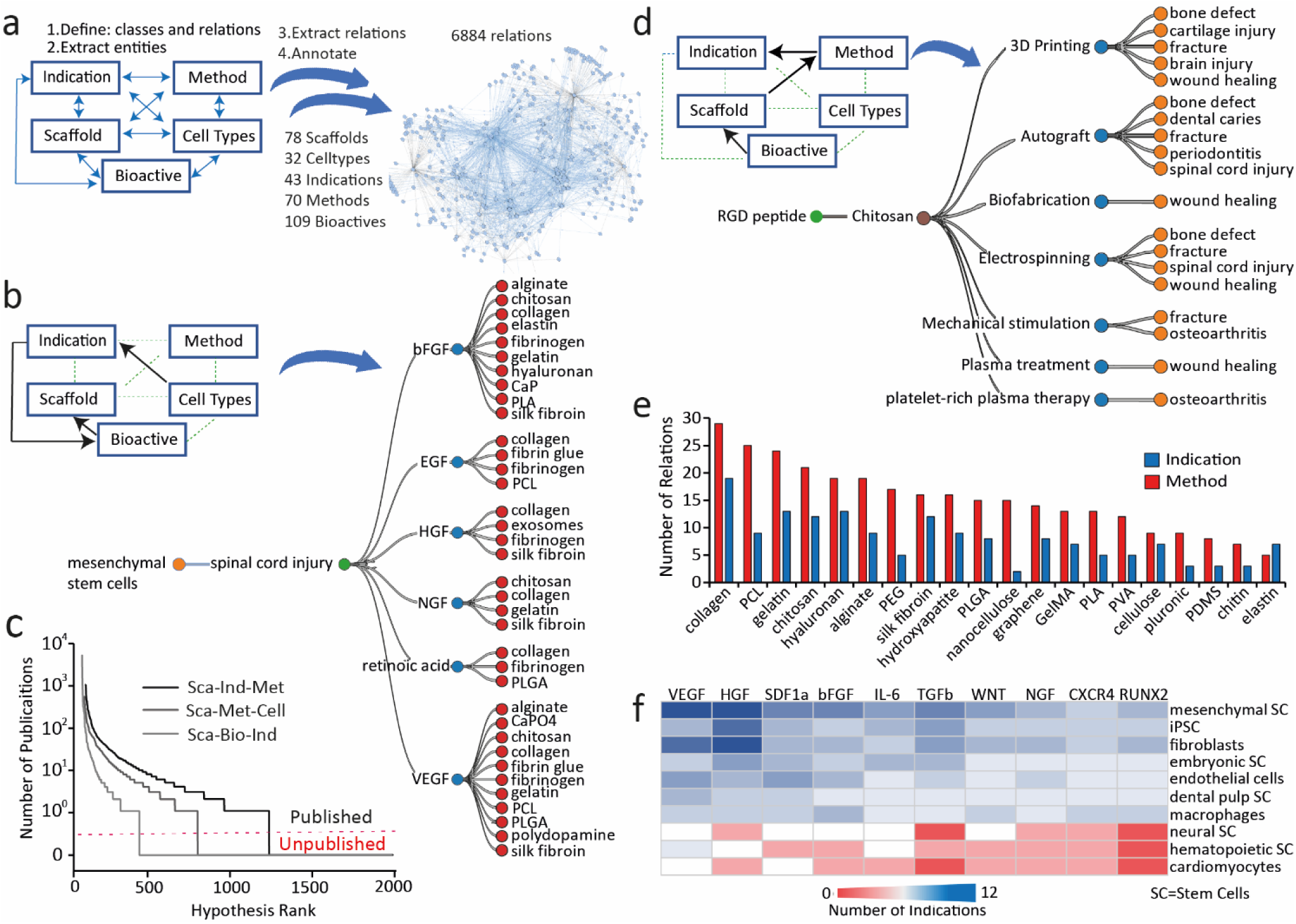
Knowledgebase for Tissue Engineering and Regeneration. **a**, knowledgebase construction scheme and details. **b,** example of hypothesis assembly in 4-entities space of sequence Celltype-Indication-Bioactive-Scaffold with filters on Celltypes (mesenchymal) and indications (spinal cord injury). **c,** 3-entities based hypotheses ranked by the number of publications in PubMed. **d**, example of hypotheses assembly starting from bioactive and ending with indication with filters on Bioactives and Scaffolds (RGD peptide and Chitosan). **e,** quantitative analysis of scaffolds relations to indications and methods. **f,** Heat map for the number of indication relations shared between Bioactives and Cell types.

After constructing a list of entities for each class (see methods), we defined the allowed relations between classes. We noted that practically all relations between all classes are highly plausible except the relation between the ‘Method-Bioactive’ classes as the vast majority methods deal with the scaffold the indication or the celltype but not the bioactive principle. We then used SPIKE-KBC app to import all published relations between entities and briefly annotated relation instances in bulk (not every single sentence) which resulted in the desired knowledgebase in about 4 hours **(Fig 3a)**. The resulting knowledge base consisted of 6884 relations between 78 scaffolds 32 cell types 43 medical indications 70 different methods and 109 bio active elements. We show a representative example **(Fig 3b)** from sequence CellType-Indication-Bioactive-Scaffold. This sequence has 3603 unique hypotheses and due to limited space, we applied filters to show a representative 10% of hypotheses space. We applied a select only filter for mesenchymal stem cells (CellType), spinal cord injury (Indication) and also filtered out randomly 10 out of 16 Bioactives and 37 out of 52 Scaffolds which yielded a total of 36 unique hypotheses which is 1% of the whole possible hypotheses space of this sequence. Of the 6 bioactive elements, two well-known growth factors, bFGF and VEGF were shown to be the most versatile and studied with 10 or more different scaffolds each. We then wished to characterize the prevalence of other hypotheses sequences in PubMed **(Fig 3c)**. We tested 3 sequences of 3 layers starting from Scaffold layer. The most published sequence was Scaffold-Indication-Method followed by Scaffold-Method-CellType and Scaffold-Bioactive-Indication. In another example we show the 4 layered sequence Bioactive-Scaffold-Method-Indication (20,000 hypotheses) with a select-in filter for RGD peptide (bioactive) and Chitosan (Scaffold) and filtered out randomly 50% of Methods and Indications to yield 19 unique hypotheses (1% from all hypotheses in the space). Following this example, the most versatile methods used with Chitosan are 3D Printing and Autograft (tissue transferred from one part of the body to another) and the most applied indication is wound healing **(Fig 3d)**. We then analyzed and ranked all the relations of the different scaffold in relations to Indications and compared to Method relations **(Fig 3e)**. We saw that most scaffolds in the top 10 are not synthetic (collagen, alginate, chitosan, hyaluronan, silk fibroin). Collagen has the greatest number of relations to both methods and indications while synthetic polymer Poly caprolactone (PCL) is ranked second in methods and 5th in indications. To overview and quantitively map the relations between Indications, Bioactives and Celltypes, we generated an unfiltered sequence Bioactive-Indication-CellTypes and analyzed the distribution of shared Indication relations to Bioactives and CellTypes (**Fig 3f**). After sorting the table, we can first observe that the most connected Celltypes are Mesenchymal Stem Cells and iPSC and Fibroblasts. It can be further seen that the VEGF and HGF -Mesenchymal Stem Cells pairs have the greatest number of shared relations. In addition, we can observe that HGF, TGFb and RUNX2 is understudied with neural stem cells and cardiomyocytes.

### Use case 2b: Cancer Surgeries and Radiotherapeutics

Since surgery and radiotherapy are the main cancer treatment modalities aside of pharmaceuticals, we moved to explore their relations with biomaterials.

The resulting knowledgebase, spanning across 440 entities and annotated with 2392 relations, was designed allow rapid hypothesis generation and exploration, aimed to evaluate more complex concepts by layering up to four classes sequentially to assemble more than 4000 different hypotheses. This construction effort was complete in about an hour, excluding entities curation. In **Fig. 4a** we show 4-class hypotheses, assembled from the class layering order: Cancer-Procedure-Tool-Biomaterial, focusing on esophageal cancer and the procedure of esophagectomy. We found 4 relations to surgical tools that in turn are related to 30 types of biomaterials – representing 30 different hypotheses. It should be noted that the sequence in which class-class relations are layered determines its reasonability. For example, the assembly of hypotheses in the sequence: Cancer-Tool-Biomaterial-Procedure, has many unreasonable relations between a procedure and a cancer as they are linked through biomaterials which are not cancer specific. On the other hand, for the space generated by Cancer-Procedure-Tools combinations, we found 68% of the hypotheses published (all terms appear together at least in one PubMed article), leading with “colorectal-colectomy-endoscope”, portraying the in-context usage of tools used in various cancer-related surgical procedures (**Fig. 4b**).

**Figure 4.**
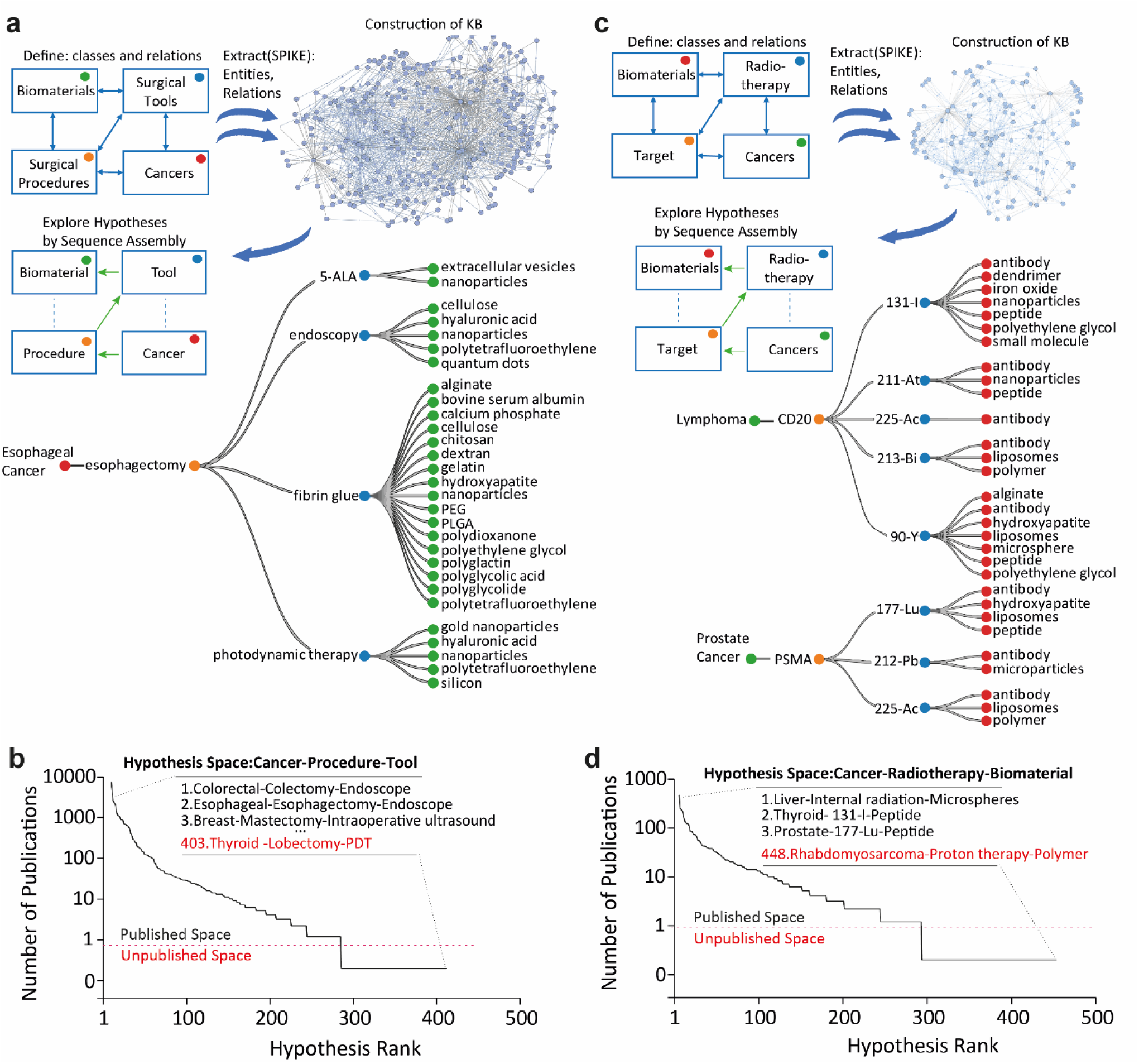
Knowledge Bases for Biomaterials in Cancer Surgeries and Radiotherapies. **a,** knowledgebase construction scheme for cancer surgeries using 4 different classes (top) and exploration in the hypotheses space using esophageal cancer as an example (bottom). **b,** 3-entities based hypotheses ranked by the number of publications in PubMed. **c,** knowledgebase construction scheme for cancer radiotherapy using 4 different classes (top) and example hypotheses assembly starting from cancer ending with biomaterials (bottom). **d,** 3-entities based hypotheses ranked by the number of publications in PubMed.

For Radiotherapies, the knowledgebase construction time was estimated at approximately a single day (including curation of entities lists) and had 120 entities and 872 relations yielding >1500 possible hypotheses. The knowledgebase was designed under the general hypothesis that different cancers are related to distinct surgical procedures or surgical instruments and are assisted by different biomaterials. The knowledgebase structure enables the user to assemble and explore localized/ precision radiotherapy hypotheses with supporting evidence. A brief analysis highlights brachytherapy as the radiotherapy having the most relations to, followed by “Intensity modulated radiotherapy”. The most common radioisotope-relations were with 131-I followed by 90-Y (**Supplementary Fig. 1b)**. Similarly, the cancers with most related radiotherapies are prostate cancer and breast cancer (**Supplementary Fig. 1c)**. Analysis of hypothesis ranking in the Cancer-Radiotherapy-Biomaterial space revealed 64% published hypotheses, leading with “Liver Cancer-Internal Radiation-Microspheres” as most published in PubMed (**Fig. 4d**).

### Use case 3: Targeted Drug Delivery Knowledgebase

Next, we sought to construct a comprehensive public knowledgebase in nanomedicine for cancer targeted drug delivery. One of the main paradigms in this field suggests that a drug (**A)**, carried by biomaterial (**B)**, can be targeted to cancer (**C)** using ligand (**D)** to bind molecular target (**E)** expressed on cell type (**F)** in the tumor microenvironment. We compiled lists of entities for each the 6 classes(A-F) – using multiple SPIKE queries (see **Table 2**) as well as databases such as DrugBank^22^, Human Protein Atlas^23^, Apta-Index^24^. We defined 10 possible relations between the classes, as shown in **Fig 5a.** The construction process resulted with 61 biomaterials, 53 Cancers, 29 cell types, 439 drugs, 219 ligands and 173 targets, connected with 6089 annotated relations which leads to >10^6^ possible hypotheses. The estimated time for construction was total of full 4-5 days spanning over 4-5 weeks. Constructing this knowledgebase took longer than the previous ones due to the carful manual annotation and name unification processes applied, which were done only briefly and in bulk in the previous knowledgebases. Another time-consuming step was curation of entity lists of biomaterials and ligands which contains many synonyms and acronyms.

**Figure 5.**
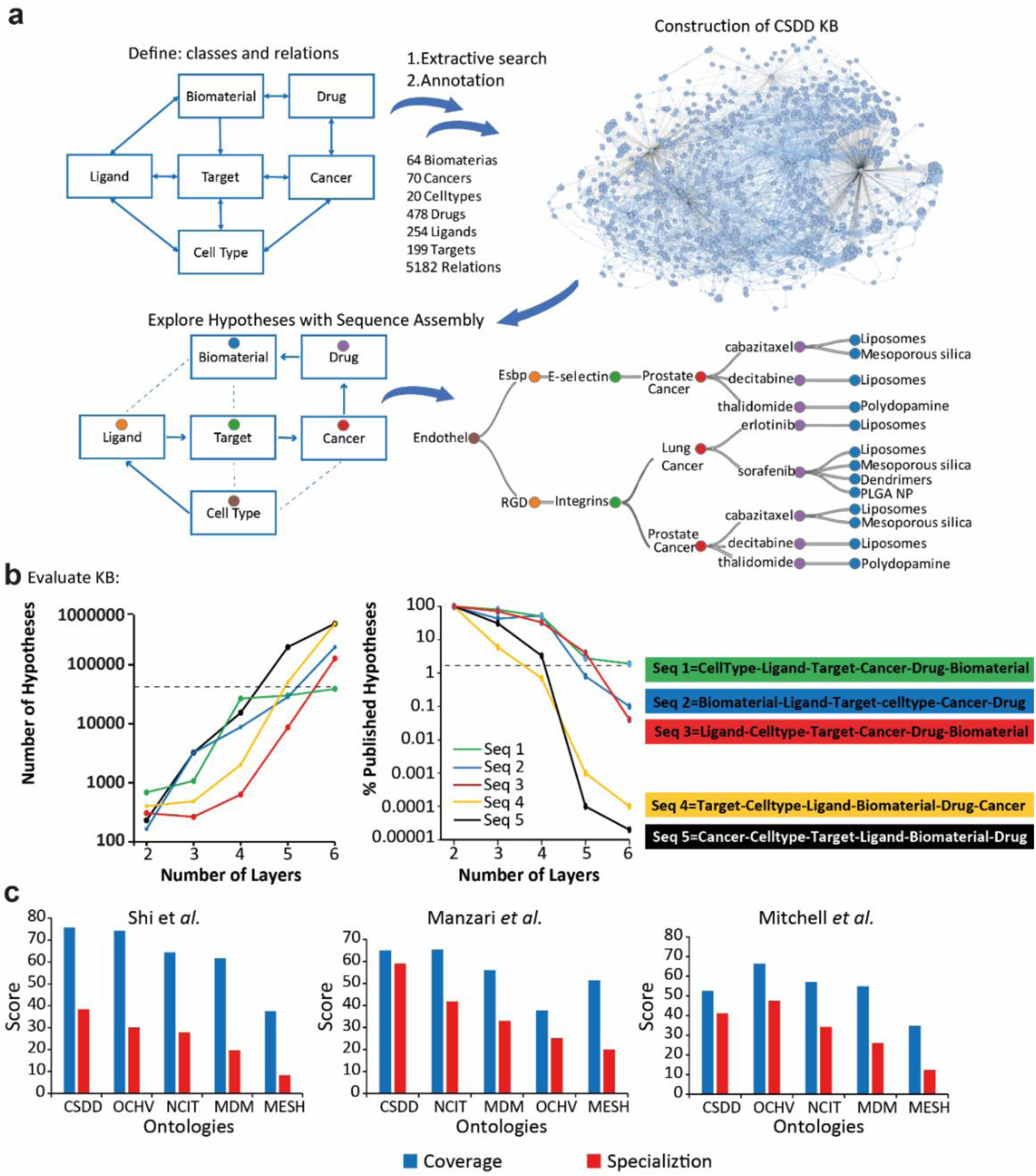
Construction of the cell specific drug delivery knowledgebase (CSDD). **a,** knowledgebase construction scheme from relation types (top). Example of exploration in the hypotheses space via sequential assembly, from biomaterials to endothelial cells (bottom). **b,** Number of possible hypotheses vs. number of layers (left); and the % of published entity combinations to the total hypotheses, arising from 5 different class layering sequences (right). **c,** Relevance of recommended ontologies to 3 leading review papers^26–28^. CSDD, Cell-Specific Drug Delivery; OCHV, Ontology of Consumer Health Vocabulary; MDM, Mapping of Drug Names and MESH 2021; NCIT, National Cancer Institute Thesaurus; MESH, Medical Subject Headings.

The estimated time for construction was total of full 4-5 days spanning over 4-5 weeks, resulting with 61 biomaterials, 53 Cancers, 29 cell types, 439 drugs, 219 ligands and 173 targets, connected with 6089 annotated relations which leads to >10^6^ possible hypotheses.

For knowledgebase exploration and hypothesis generation, we show an example for targeting endothelial cells in prostate or lung cancers is shown in **Fig. 5a** (bottom). Of the 26 possible class layering orders, based on the available annotated relations, the 4 most promising sequences were evaluated. Critical assessment revealed that arbitrary class layering orders results with an overwhelming number of possible entity combinations (**Fig. 5b** left), while having very few correlating references in the PubMed corpus (low % hits). In contrary, sequence #1 (cell type ► ligand ► target ► cancer ► drug ► biomaterial) consistently yielded the most hits with 2.5% of the suggested combinations represented in PubMed, while all the other sequences fall below 0.01% (**Fig. 5b** right). As this knowledgebase was aimed to be comprehensive and accurate, we evaluated the knowledgebase’s coverage and specificity relevance to its stated discipline of targeted drug delivery. In **Fig. 5c** we show the average coverage and specialization scores of the CSDD knowledgebase and 4 other leading ontologies, calculated by the NCBO Ontology Recommender^25^, with respect to entities mined from selected review articles (Supplementary **Table S1**)^26–28^. The coverage score, representing the extent to what the ontology covers the input data, similarly to the level of specialization of the ontology to the domain of the input data, were both comparable or superior to other ontologies.

Following the CSDD knowledgebase overview and quality measures, we move to its meta-analysis. To direct the optimal class layering order, we compared the prevalence in PubMed of the combinations of entities, generated by different class layering sequences. In the Cell Type-Cancer-Target space, 82% of hypotheses were published, with leading hypothesis “Cancer cells-Breast Cancer-HER2”, followed by “B-cell-Lymphoma-CD20” (**Fig. 6a**). Indeed, HER2 and CD20 targeting is one of the most used examples in precision nanomedicine^29, 30^. In contrast, the combination “Endothelial cells-ANXA1-Gastric Cancer”, is an example of hypothesis that is unpublished, even though every pairwise relation within it had published evidence. In the Biomaterial-Drug-Cancer space, 71% of hypotheses are published. The most frequent hypothesis is “Liposomes-Doxorubicin-Breast cancer” followed by “PEG-Doxorubicin-Breast cancer” **(Fig. 6b)**. Again, we saw that the most common hypotheses are ones including the most published cancer (breast cancer), drug (doxorubicin) and biomaterials (liposomes). In contrast, “Chitosan-Rapamycin-Melanoma” contains multiple published pair relations even though the explicit combination is unpublished. The same analysis, applied for Cancer-Target-Biomaterial revealed “Hyaluronan-CD44-Breast Cancer” as the leading hypothesis followed by “Liposomes-EGFR-Breast Cancer” (**Supplementary Fig. 2a)** and ‘Doxorubicin-Hyaluronan-CD44’ for Drug-Biomaterial-Target (**Supplementary Fig. 2b**). Next, we analyzed specific entities and their prevalence in relations in different contexts. We found that the targets with most biomaterial-drug relations are the Endothelial Growth Factor-Receptor (EGFR) and CD44 (**Fig. 6c**). In the context of targeting biomaterials, the most popular ligands are folic acid and the cell penetrating peptide, R8 (**Fig. 6d**). Next, as cancers and cell types are highly associated, we assessed the number of different targeting candidate per each cancer and cell type. As can be seen in **Fig. 6e**, multiple targets are shared between different cancers and cell types. While Esophageal cancer targeting was found only to a few ’Cancer cells’, ‘Fibroblasts’ and Cancer stem cells’ targets, Breast cancer active targeting is suggested by many hypotheses, involving multiple targets for different cell types: 21 different targets for ‘Cancer cells’, 15 to ‘Endothelial cell’ etc.

**Fig. 6.**
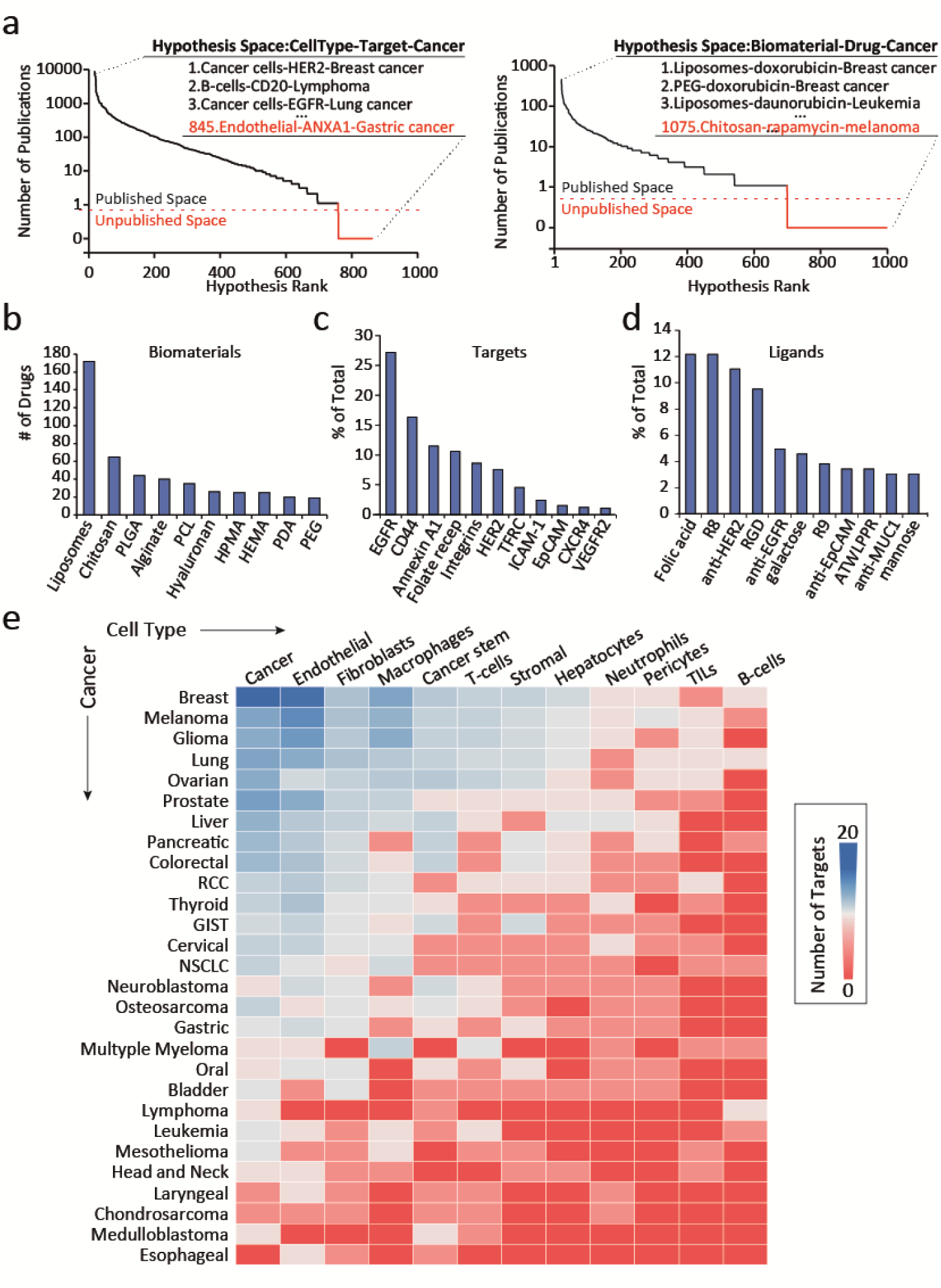
Meta-analysis of the CSDD knowledgebase. **a,** 3-class entity combinations, derived from two different general hypotheses, ranked by the number of publications in PubMed. **b,** Prevalence of biomaterials by the number of drug relations. **c,** Relative distribution of targets in combinations with both cancers and biomaterials. **d,** Relative distribution of targeting ligands relations to both targets and biomaterials. **e,** Heat map for the number of targets relations shared between cancer and cell types.

## Discussion

Existing approaches to biomedical knowledgebase construction require significant specialization and expertise, and a significant time investment. On the one hand, there are biomedical ontologies which are constructed by professional Biocurators. These are highly accurate, but can take years to construct and maintain^7, 8^. On the other hand, there are knowledgebases which are automatically constructed through the use of ML models^12, 31^. While ML models greatly simplify the task of keeping the knowledgebases up to date, their initial creation requires significant expertise in NLP, ML, and ML productization, as well as significant time investments in data annotation, evaluation, and model tuning. In practice, both biocuration and automatic knowledgebase construction involve inter-lab collaborations and are typically out of reach for individual researchers or small labs. As a consequence, existing knowledgebases tend to be focused on relation types that are expected to be of interest to a wide scientific community, thus leaving a wide “tail” of relations which are not covered by any knowledgebase.

We demonstrated that by using extractive search, we can significantly lower the barrier of entry for knowledgebase construction, allowing individual researchers or small teams to rapidly construct personalized, targeted knowledgebases that are tailored to their research interests and are anchored in the literature. The effort required to create such knowledgebases depends on the researchers’ goals and their expected level of coverage and accuracy, and can range from hours to weeks.

In the results section of this paper, we surveyed different knowledgebases which manifest the above-mentioned tradeoffs. The knowledgebases were constructed by researchers in a biomedical engineering lab specializing in targeted drug delivery and cancer research. The constructed knowledgebases reflect some of the main research topics in biomedicine: the cancer frontiers knowledgebase is designed to identify recent trends and challenges in cancer research. It was created by a single researcher in a couple of hours, by aggregating the results of a few extractive queries without validation. The knowledgebase can be easily kept up to date by re-applying these queries, and since the knowledgebase is for personal use, occasional errors or omissions are not a significant concern. The tissue engineering and regeneration, cancer surgery and radiotherapy knowledgebases were each created in less than a day by applying sporadic validation to the results of extractive queries. These knowledgebases were used to support rapid hypothesis exploration and generation as discussed in the Results section. For this purpose, non-prohibitive amounts of noise were acceptable: the knowledgebases are used to surface promising hypotheses, but each hypothesis is directly linked to the literature, and can subsequently be reviewed by researchers who use their experience to focus on useful ones and eliminate those based on incorrect connections. Finally, the CSDD knowledgebase was designed to be a comprehensive and up to date source of structured knowledge for the wider drug-delivery research community. To maintain the levels of precision expected from a public resource, it was constructed by 3 annotators over the course of a few weeks and all curated facts were manually validated.

In surveying these knowledgebases, we demonstrate that knowledgebase construction need not be an effort-intensive process. By applying extractive queries and varying degrees of manual validation, researchers can rapidly create knowledgebases to accelerate their own research and to benefit the larger community.

## Conclusions

We propose a fully functional human-machine hybrid method for rapid construction of both personalized and public knowledgebases in biomedicine and demonstrate the construction and evaluation of five different knowledgebases. The resulting knowledgebases facilitate hypotheses generation and exploration backed with evidence from the literature. We provide the infrastructure, the tools and tutorials for a broad use of diverse biomedical researchers with no background in text mining and NLP. The potential of generating plausible hypotheses that are yet unpublished can enable accelerated discovery in multiple fields of biomedicine.

## Methods

The presented workflow repeats, with minor modification, in all of the reported knowledgebases. In each case, we commence with project definition and the implementation of extractive queries to (i) obtain lists of biomedical entities, and (ii) capture relations between these entities, adding these relations and their supporting evidence to the knowledgebase. The lists and queries used for the construction of the different knowledgebases are detailed below, and are also available in the supplementary information along with instructions on how to easily load them to SPIKE-KBC to reproduce our findings.

### Construction of the Challenges in Cancer Research Knowledgebase

#### Project definition

This case demonstrates a literature review task of capturing unknown entities and concepts from intangible classes. We aimed to construct a knowledgebase of recent challenges in cancer research using three conceptual classes: ‘controversial’, ‘still unknown’ and ‘unmet need’.

#### Generation of entity lists

Independent of any prior hypothesis on what concepts might be ‘controversial’, we used SPIKE over PubMed search engine to mine concepts into *de novo* name lists, applying structured (syntactic) queries^15^. For example, to retrieve controversial findings we used the query “<>:something $remains/still/is $controversial”, restricted to papers that have words pertaining to different cancer types in their abstract and filtered to the last three years to reflect only contemporary topics. In this query, a colon (‘:’) was used as captures for things that are controversial, ‘<>’ to expand the captured string to possibly more than one word, and ‘$’ as an anchor, requiring one of the specified following words to appear in the captured sentence. Similar queries were used to capture possible ‘unmet needs’ and ‘things that are unknown’ (see **Table 1a**) The terms were saved as lists within the SPIKE system for later use.

**Table 1a:** Extractive search queries used to mine entities based on their class, to be used in the Challenges in Cancer Research knowledgebase.

| Query Type | Filters | Sought Class | Query | # of Captured Entities |
| --- | --- | --- | --- | --- |
| Structure | Year field:2018 to 2022,<br>Abstract: "cancer" | Controversial concepts | <>:something \$is \$controversial | 802 |
| | | Controversial concepts | <>:something \$remains \$controversial | 47 |
| | | Controversial concepts | <>:something \$still \$controversial | 3 |
| | | Controversial concepts | the \$controversy \$of <>:something | 16 |
| | | Unknown concepts | <>:something \$remains \$unknown | 14 |
| | | Unknown concepts | <>:something \$is \$unknown | 1174 |
| | | Unmet needs | <>:something \$is \$unmet \$need | 86 |
| | | Unmet needs | <>:something \$remains \$unmet \$need | 16 |
| | | Unmet needs | \$unmet \$need \$is <>:something | 10 |
| | | Unmet needs | the \$unmet \$need \$of <>:something | 21 |

#### Mining candidate relation instances

To match various cancers to their corresponding ‘challenging’ concepts, citations having two sought words appear in the same sentence, were mined from the PubMed corpus. In the SPIKE search engine, a list of 47 cancer types, based on the National Cancer Institute’s list of cancer types^32^, was matched to one of the previously formed lists using the Boolean extractive query ‘arg1:{listA} arg2:{listB}’. In this query, ‘arg1’ and ‘arg2’ are arbitrary argument names and the braces (‘{}’) act as placeholder to enable the automatic, iterative search of all words within a pre-saved concept list with the corresponding name (e.g., ‘arg1:{Unmet Needs} arg2:{Cancers}’). By applying this search, SPIKE retrieved citations that capture matching pairs of specific cancer type-controversial entities in the same context, tagging the specified entities within the sentences (see Table 1b).

**Table 1b:**
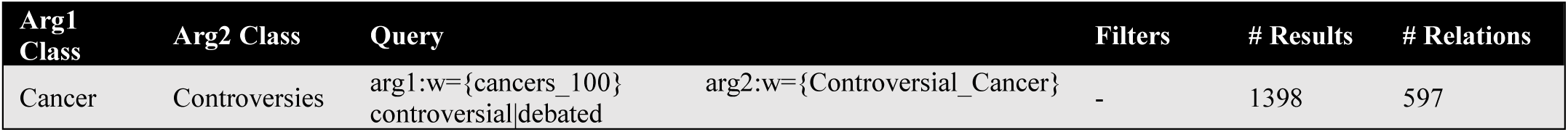
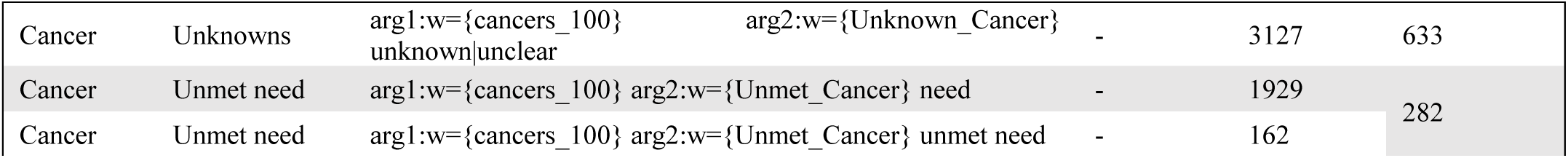
Queries used for the capture of relation instances for the Challenges in Cancer Research knowledgebase. The number of results include redundant instances of identical relations, before their unification as the net number of approved relations.

### Validation and annotation of relation instances

The captured citations were exported from SPIKE by copying the query’s SPIKE URL and imported into the SPIKE-KBC app. For each of the 3 conceptual categories, a separate relation was added. The captured sentences, grouped by and tagged with the specific entity combination as ‘arg1’ and ‘arg2’, were individually reviewed by the annotator to be appraised as appropriate relation instances. Per each entity-entity combination, the annotator decides whether a certain relation instance (the annotation) complies with the defined relation, and whether it should be included in the knowledgebase, given three options: Approve, where both decisions are positive, which results with the relation instance included in the knowledgebase with the citation as evidence; Alternatively, Reject opts to discard the relation completely from the knowledgebase, with all the corelating instances. Lastly, the Delete opts to discard an irrelevant instance that fails to support the specified relation, though unlike Reject, it does not reject the relation itself, which may be supported by further evidence for annotation. In this case, since this is an explorative knowledgebase, and in order to avoid bias caused by the annotator’s prior knowledge, all relations were approved in bulk, except boldly out-of-context entities or relations. The proposed instances are ranked by default such that frequently occurring candidates are shown before rare ones. However, reversing this order highlighted undiscovered or rare connections, was found useful in this context. For this knowledgebase and all that followed, unless stated otherwise, a minimal threshold of instances per entity pair was not applied. Consequently, a single reference was adequate to establish a connection.

#### knowledgebase Analysis

The visualization module was used to review and analyze the resulted knowledgebase. We displayed each of the two connected classes (e.g., cancer<->unmet needs) and downloaded a .csv file of all the two entity pairs, allowing to compare the total number of relations per cancer. Reversibly, we counted the total number of relations per concept, highlighting the concepts having the most relations to (e.g., ‘what are the most unmet needs in cancer?’).

### Construction of Tissue Engineering and Regeneration Knowledgebase

#### Project Definition

The field of tissue engineering and regeneration is one of the major disciplines in biomedical engineering and could be a great example for rapid knowledgebase construction. We set to define the main hypothesis in this field with a combinatorial relation of classes. Tissue engineering is defined as the use of various cells, material engineering methods and biochemical factors to restore, maintain, improve, or replace biological tissues^21^. We took this simple definition and reconstructed it to the following combinatorial hypothesis of entity classes: A certain medical condition **A** can be treated with a scaffold made from material **B** which can support the growth of cell type **C** by combining bioactive element **D** (growth factors, hormones). We also noted that there many different methods to combine the cells with scaffolds and how to introduce the scaffold to the damaged tissue and for that purpose we added another class Method **E.** After defining each class, we defined the allowed relations between classes. We noted that practically all relations between all classes are highly plausible except the relation between the ‘Method-Bioactive’ classes as the vast majority methods deal with the scaffold the indication or the Cell Type but not the bioactive principle.

#### Generation of Entity Lists

After defining the required classes, we moved to extract entities for each class. For that purpose, we used the SPIKE engine and a series of structured queries to produce lists of entities (see methods). In brief we used extractive queries such as ‘<>:something $is $method|technique’ and applied a filter in the abstract: ‘tissue engineering|regeneration’. This approach mostly captured words that are types of methods used in tissue engineering. They were then manually selected as appropriate entities from the displayed sentences (see **Table 2a**).

**Table 2a:**
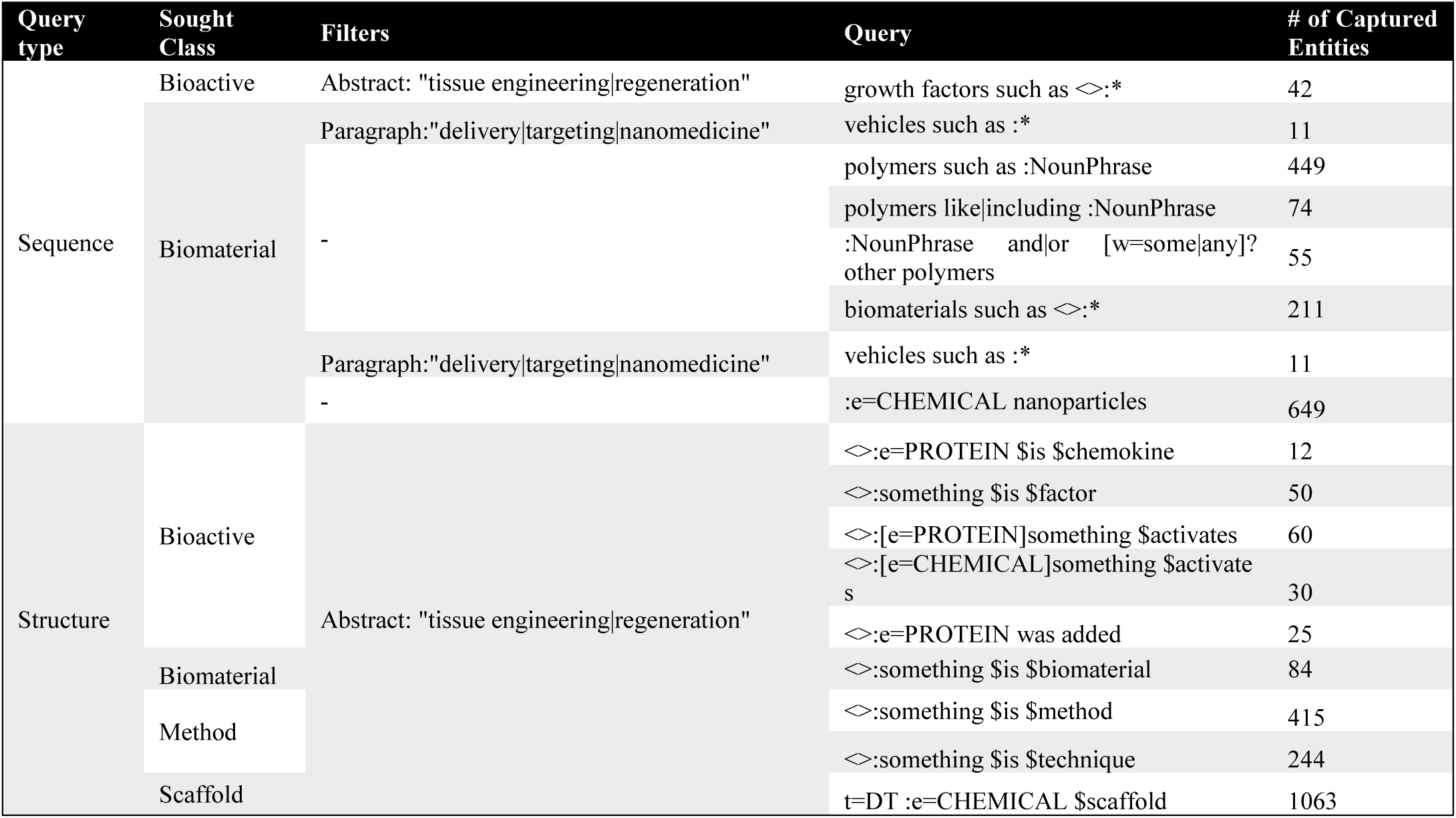
Extractive queries used to capture entities and compile entity lists in the Tissue Engineering and Regeneration knowledgebase. t=DT is a tag for a determiner such as ‘the’. ‘e=’ directs the us of a term list, buit-in SPIKE.

| Query type | Sought Class | Filters | Query | # of Captured Entities |
| --- | --- | --- | --- | --- |
| Sequence | Bioactive | Abstract: "tissue engineering\|regeneration" | growth factors such as <>:* | 42 |
|  | Biomaterial | Paragraph:"delivery\|targeting\|nanomedicine" | vehicles such as :* | 11 |
|  |  | - | polymers such as :NounPhrase | 449 |
|  |  |  | polymers like\|including :NounPhrase | 74 |
|  |  |  | :NounPhrase and/or [w=some\|any]? other polymers | 55 |
|  |  |  | biomaterials such as <>:* | 211 |
|  |  | Paragraph:"delivery\|targeting\|nanomedicine" | vehicles such as :* | 11 |
|  |  | - | :e=CHEMICAL nanoparticles | 649 |
| Structure | Bioactive | Abstract: "tissue engineering\|regeneration" | <>:e=PROTEIN \$is \$chemokine | 12 |
| | | | <>:something \$is \$factor | 50 |
| | | | <>:[e=PROTEIN]something \$activates | 60 |
| | | | <>:[e=CHEMICAL]something \$activate s | 30 |
|  |  |  | <>:e=PROTEIN was added | 25 |
| | Biomaterial | | <>:something \$is \$biomaterial | 84 |
| | Method | | <>:something \$is \$method | 415 |
| | | | <>:something \$is \$technique | 244 |
| | Scaffold | | t=DT :e=CHEMICAL \$scaffold | 1063 |

#### Mining candidate relation instances

Standard Boolean queries, as described above, were used to mine instances where two specific entities appear in the same context (see **Table 2b**). The captured citation, tagged with the entity pairs, were imported to SPIKE-KBC.

**Table 2b:**
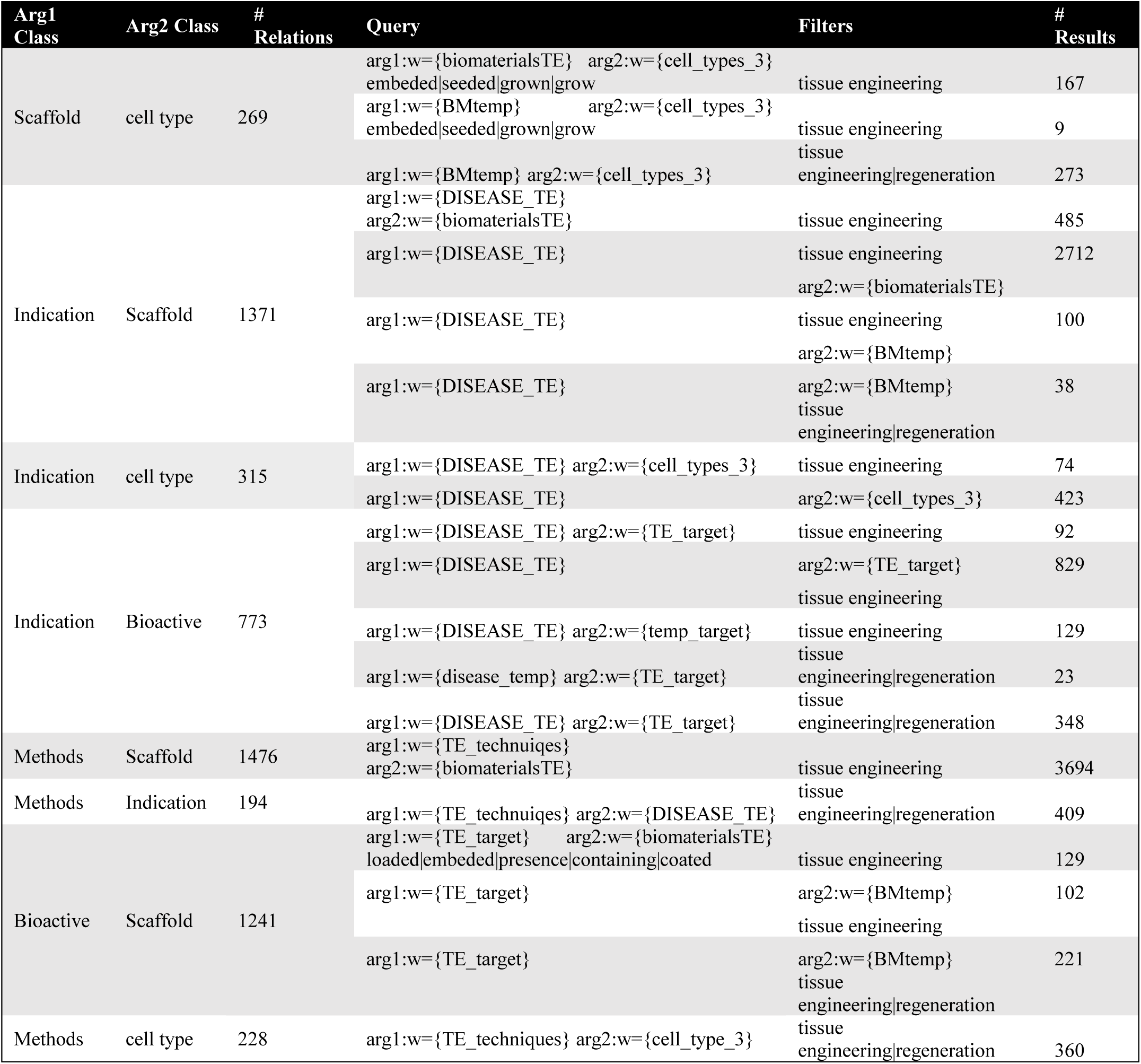
Queries used for the capture of relation instances for the Tissue Engineering and Regeneration knowledgebase. All queries were filtered to include ‘Tissue engineering’ or ‘Tissue Regeneration’ in the Abstract.

#### Validation and annotation of relation instances

As indicated above, the SPIKE-KBC app was used to review all the mined relations between entities. The relation instances were briefly reviewed and annotated in bulk (not every single sentence, but per entity pair).

#### Knowledgebase Analysis

To explore the knowledgebase dedicated to Tissue Engineering and Regeneration, we applied similar analysis to the CSDD; per each multi-class visualization and applied filters, a comma-separated values (.CSV) file was downloaded. The prevalence of an entity to be related to other of different class was enumerated using a pivot table, rendering the count of entities of class A as values for the second class as categories. Similarly, a two-variable matrix of ‘Bioactives’ and ‘Cell Types’ was rendered with the shared ‘Indications’ as values. To evaluate the class layering order (sequence) which produce entity combinations that are most published on PubMed, we tested different 3 or 4-layers sequences. For each of the specific 3 or 4 entities combinations, the number of search results in sought in PubMed and was summed.

### Construction of Cancer Surgeries and Radiotherapeutics Knowledgebases

#### Project Definition

The driving hypothesis for **Cancer Surgeries** knowledgebase is that specific malignancies correspond with certain surgical procedures, in which surgical instrument and biomaterials are used. Accordingly, 6 hypothetic relations were phrased, involving cancers, surgical procedures, surgical tools and their relations to biomaterials (see **Supplementary Video 1)**.

The same approach was used for the construction of a similar, complementary knowledgebase on **Targeted Cancer Radiotherapies**, consisting of 4 classes of interest: Cancer, Target, Radiotherapeutic agent, and Biomaterial. The 4 defined relations are based on the general paradigm of targeted radiotherapeutics, according to a radiotherapeutic agent can be administrated in cancer treatment using targeted biomaterials. We note that diagnostics were omitted from this knowledgebase.

#### Generation of Entity Lists

To establish lists of surgical procedures and surgical tools, we used a similar approach to the described above, and used Sequence (token) queries to mine biomaterials, surgical procedures etc. based on expected use in sentences. Special syntax was used to direct the search of specific parts of speech (see **Table 4a**).

**Table 4a:**
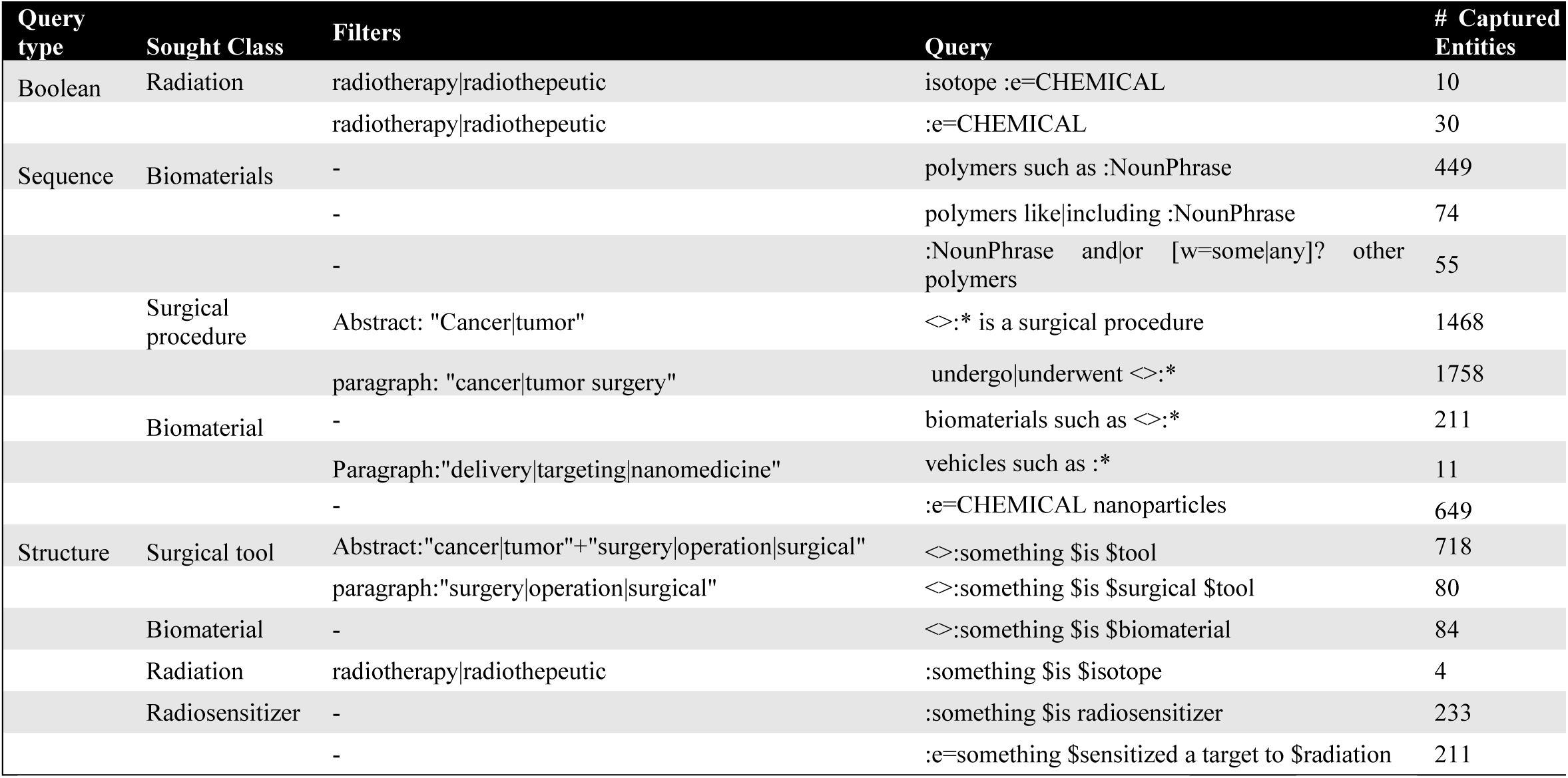
**Extractive queries used to capture entities and compile entity lists in the Cancer surgeries and radiotherapy knowledgebases.** ‘NounPhrase’ is shorthand for “(([t=DT]? [t=/JJ.*/]* [t=/NN.*/]+))” directing a determiner, followed by 0 or more adjectives followed by 1 or more nouns). Asterix (*) is a wild card to match any single word. lists previously mined are not shown.

**Mining candidate relation instances:** standard Boolean queries were used to capture relation instances between the defined classes, as demonstrated above (see **Table 4b).**

**Table 4b:**
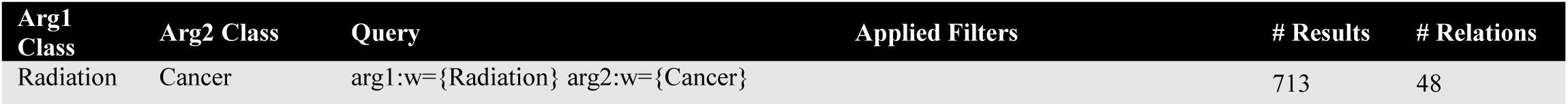
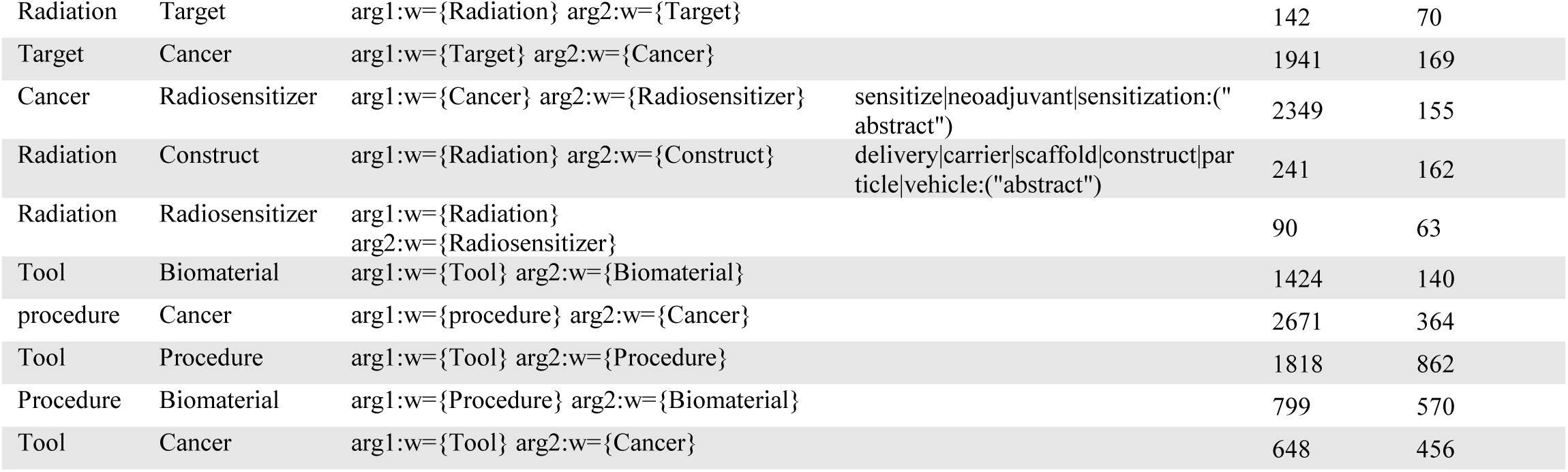
Queries used for the capture of relation instances for the cancer surgeries and radiotherapy knowledgebases.

#### Validation and annotation of relation instances

The results of the queries were imported from SPIKE to SPIKE-KBC as described above. The annotation process was done in bulk and was not supervised under specific criteria. The process was recorded and explained in real time for the purpose of demonstration and can be viewed here: https://www.youtube.com/watch?v=zRxnNfdQNF0&t=25s

#### Knowledgebase analysis

Different class layering sequences were visualized to generate and explore different hypotheses in radiotherapy or cancer surgery. We evaluated the published space derived of each class-layering order, as thoroughly described above. 3 sets of 3-class combinations and 3 sets of 4-class combinations were compared, each generating unique entity combinations. The sets were searched in PubMed and the overall number of results was counted.

### Construction of Cell Specific Drug Delivery Knowledgebase

#### Project Definition

To construct a comprehensive public knowledgebase in nanomedicine for cell specific targeted drug delivery, a generally phrased hypothesis that conveys the topic’s main paradigm was laid to define the knowledgebases outlines, as follows: A drug (A), carried by biomaterial (B), can be targeted to cancer (C) using ligand (D) to bind molecular targets (E) expressed on cell type (F) in the tumor microenvironment. Of the 15 possible relations between the classes, 9 of interest were defined, comprising of two entity classes and their appropriate logical relation, such as “Drug” and “Disease”, and the string “is_used_to_treat” as their relation. While some relations are obvious (e.g., ligand-target), the reasoning for the inclusion or omission of certain relations was done according to the general hypothesis’ logic: Biomaterials, aside from their role as scaffolds or carriers for the loading of a therapeutic agents or a targeting molecules (ligands), often convey useful intrinsic characteristics on their own (e.g., protein binding, immune cell evasion etc.). Hence, the relations between specific targets, ligands, drugs and biomaterials may express meaningful, specific interactions. Additionally, as various cell types are known to be involved in the tumor microenvironment and are considered as possible targets in some treatment modalities (such as, the cytokine targeting of immune cells), this relation was included. Lastly, the use of binding molecule to target or label certain cell types is often described in the literature without relating to a specific target, therefore the ligand-cell type relation was included.

**Generation of Entity Lists** was performed by applying a hybrid approach to maximize the knowledgebase’ coverage – partially mining entity lists *de novo* using SPIKE extractive search, as previously described (**Table 3a**), and partially manually mining name lists from complementary databases and ontologies (**Table 3b**). In SPIKE, peptide-based targeting molecules (ligands) were mined using a case-sensitive query, to capture 7 to 14 letters-long strings written in upper case – a formatting almost exclusive for single-letter notation of amino acids. We listed all human Clusters of Differentiation (CDs) as possible targets, to be matched with corresponding cell types and cancer^33^. respectively, a reciprocal, hypothetic list of ligands, was created by adding the string ‘anti-‘ before all CDs, to facilitate the mining of relations to matching ligands (e.g., ‘CD40’ and ‘anti-CD40’ antibody).

**Table 3a:**
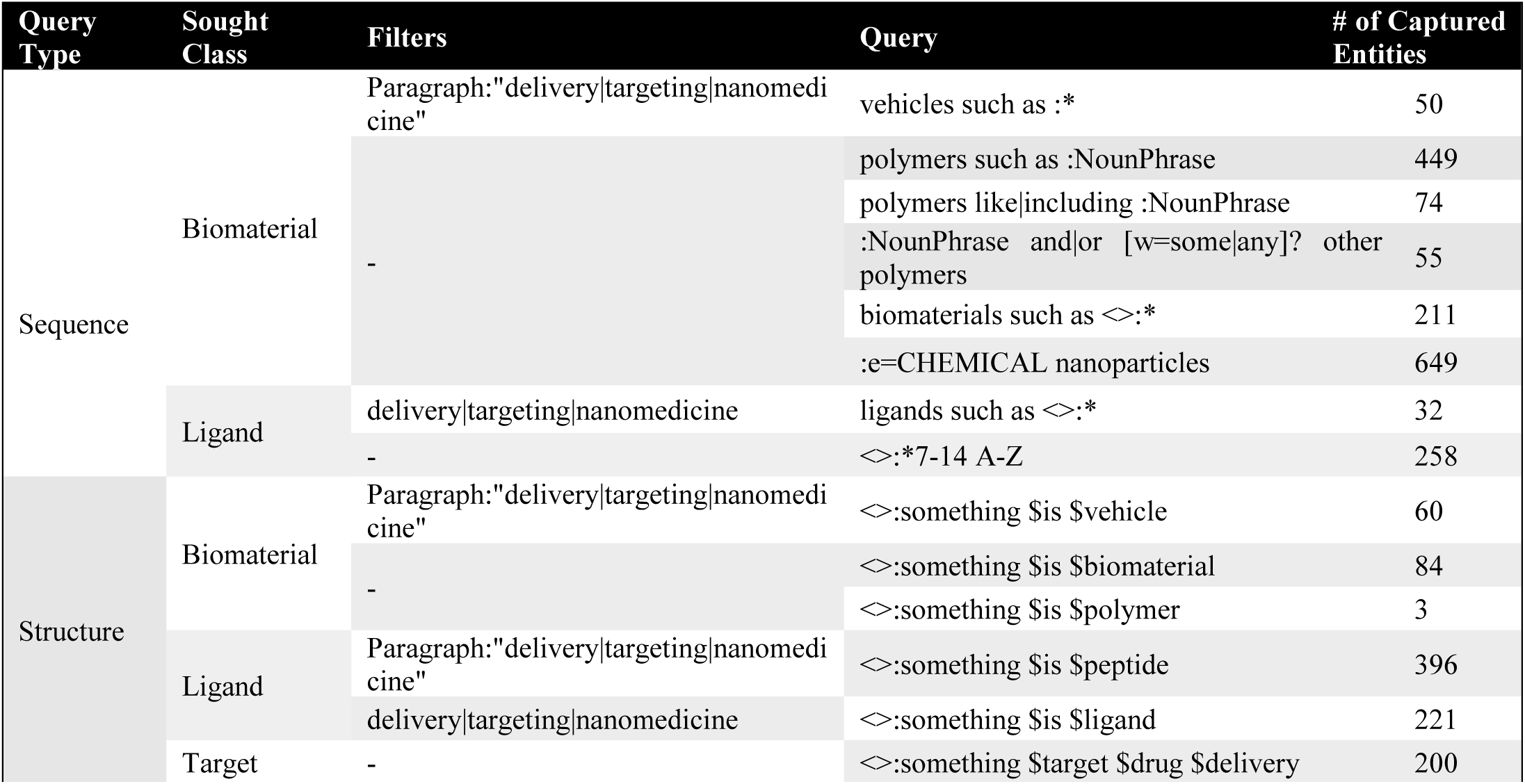
Extractive queries used to capture entities and compile entity lists in the Targeted Drug Delivery knowledgebase. ‘NounPhrase’ is shorthand for “(([t=DT]? [t=/JJ.*/]* [t=/NN.*/]+))” directing a determiner, followed by 0 or more adjectives followed by 1 or more nouns). Asterix (*) is a wild card to match any single word.

**Table 3b:** Complementary sources for acquirement of entities for the Targeted Drug Delivery knowledgebase.

| Database | Class | number of entities | Reference |
| --- | --- | --- | --- |
| DrugBank | Drugs | 910 | 27 |
| Apta-Index | Ligands (aptamers) | 87 | 29 |
| HCDM | Targets (CD) | 400 | 28 |
| Uniprot | Targets | 392 | 30 |

Additionally, aptamer-based ligands were mined from Apta-Index^24^, DrugBank^22^ was used to retrieve a list of drugs, and the Uniprot^34^ database was a source for targets. A complementary list of biomaterials and drugs were retrieved from the author’s own inventory.

#### Mining candidate relation instances

Assuming the existence of a relation between two specific entities, it will have evidence in the published literature. Hence, extractive queries were used to find instances of their co-occurrence in text, to possibly support the sought logical relation. Similarly to the described method above, 2 lists of discrete entity classes, added to SPIKE in the previous step, were used in the pairwise Boolean query ‘arg1:{ligand_list} arg2:{target_list}’. In this query, ‘arg1’ and ‘arg2’ were used as placeholders to enable the automatic, iterative search of pairwise combinations of entities from each list, and the retrieval of matching citations within the PubMed corpus, having the two words appear in the same sentence. Some queries, as detailed in **Table 3c**, include domain specific terms which are required to be in the same sentence, paragraph or section, restricted by specific titles or journals. The resulting relation instances, tagging the specified entities within the sentences as possible relation instances, were exported from SPIKE by copying the query’s SPIKE URL and imported into the SPIKE-KBC annotation module.

**Table 3c:**
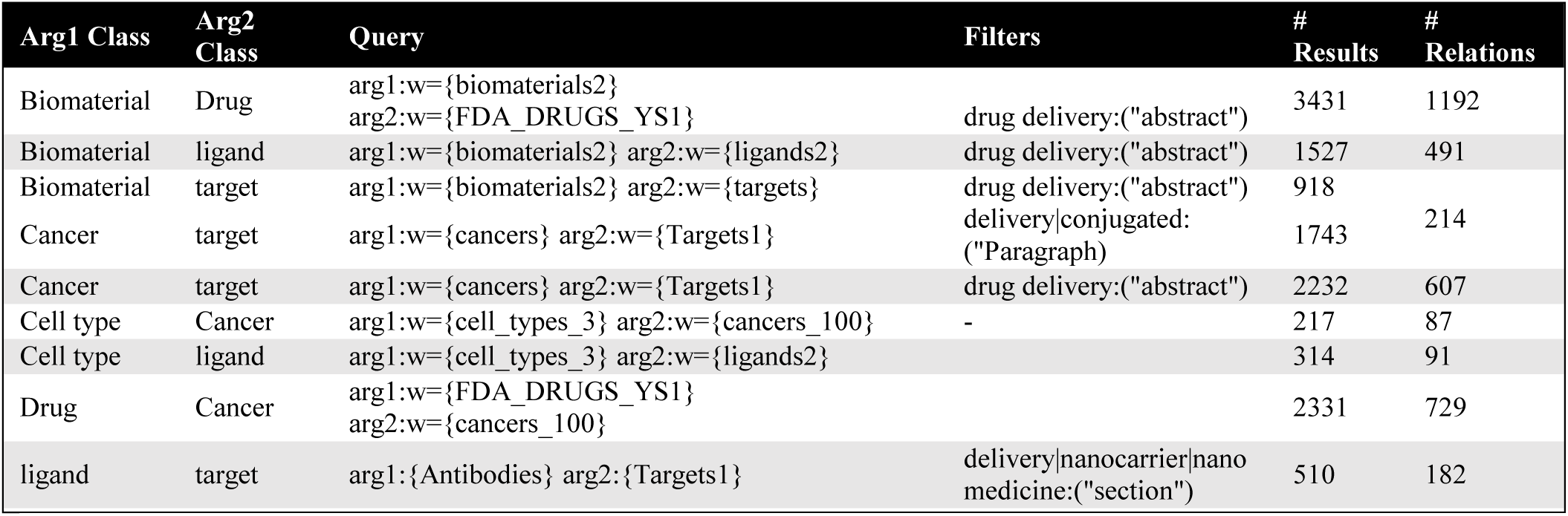
Queries used for the capture of relation instances for the Targeted Drug Delivery knowledgebase.

#### Results Validation and Annotation

As described above, for each case of class-class relations, the appropriate two-argument queries were imported and loaded into the annotation module. a team of 3 annotators reviewed the results. For this use, once approving or rejecting an instance, additional instances of the same entity pair were mostly skipped by option, thus the number of instances in the final knowledgebase are not proportional to the original number of cited instances given by the search query. **Post Processing:** In all knowledgebases, some relation instances were retrospectively reviewed and individually removed, only if they included a capture error (e.g., marking the string ‘practical’ as a Target).In the CSDD knowledgebase, as this knowledgebase is in the Authors’ own specialty, post-processing was practiced to removed context-irrelevant or boldly false relations, such as ‘anti-CD47’ (Ligand) related to ‘CD4’ (Target). The manual name unification module was applied as a quality measure and to increase the knowledgebases’ coherency and integrity. According to the annotator’s discretion, unwanted duplicates of separate entities were merged into a single entity, to override aliases, acronyms, alternative delimiters, letter capitalization, or use of abbreviations.

#### Knowledgebase Analysis

To shed light on the space of knowledge and prevalence of different entities in a defined context, different visualizations, such as all the Drug-Biomaterial relations, were downloaded as a .csv file and were transformed to a pivot table using Microsoft Excel. The prevalence of the use of the different targets or ligands, as % of total, was compared by counting, per specific entity, the number of relations exist to other entities. Similarly, by layering 3 classes and downloading all the specific paths, the number of relations shared between two classes to a third one was counted, having two classes as categories and the third as values. The resulting matrix was colored via conditional formatting, highlighting the over or under-shared specific entities.

#### Critical Evaluation

Bearing in mind that the multi-class hypotheses are constituted from discrete, two-class relation instances tailored together, the order of their layering affects the logics of the hypotheses. To evaluate which sequence is more reasonable, we systematically compared all entity combinations (hypotheses) arising from a specific class layering sequence (Set A) to fraction of entity combinations existing within the same text in PubMed (Set B). First, the crude size of each set (|*A*|) was compared according to different layering orders, starting with 2 layers and up to 6. Secondly, the intersection *B* ∩ *A* = *A*’ are entity combinations arising from a given class sequence, also appearing within discrete texts in PubMed. As A’ is a subset of A (*A* ∩ *A*’), the relative hit percentages were compared (%*hits* = 100 x |*A*|/*A*|’), where |A| is the number of entity combinations sought for in Pubmed, and |*A*|’ is the number of times such entity combinations appear together in PubMed papers (number of ‘hits’). Accordingly, if % hits = 100, all combinations are found within PubMed texts; whereas if % hits = 0, none of the combinations could be found within PubMed texts. This was calculated per each sequence, one layer at a time, until 6 classes were used in combination.

Lastly, we sought to evaluate whether the knowledgebases meets its goal of representing the current state of knowledge, with respect to comprehensive reviews in the field. The CSDD knowledgebase was exported in OWL format and imported to Bioportal^20^, where Bioportal’s ontology recommender^25^ was used to compare the CSDD knowledgebases coverage and specialization with other published ontologies, with respect to highly cited review articles in the corresponding field^26–28^. Biomedical entities were manually mined from the 3 texts and were used as keywords for calculating ontology recommendations (see **supplementary table s1**).

### Data availability

The SPIKE-KBC system and the knowledgebases reported in this work are publicly available at https://spike-kbc.apps.allenai.org.

Complete tutorial on how to construct and explore knowledgebases using cancer surgeries as an example: https://www.youtube.com/watch?v=zRxnNfdQNF0&t=25s

The source code for SPIKE-KBC as well as the lists and queries used for the construction of the described knowledgebases are available in the supplementary information and will be made publicly available under the open source Apache license.

### Code availability

The code will be posted as Github repository after review of the code upon publication.

## Competing Interests Statement

The authors declare that there are no conflicts of interests.

## Acknowledgements

We thank Aptagen LLC (Jacobus, PA) for providing access to the Aptamer Index; Amotz Taub-Tabib for annotating the CSDD knowledgebase. YS would like to thank funding from ISF grant #901/91.

## Author Contributions

YS and HT conceived the protocol and platform. SLW and YS constructed all the knowledgebases including annotations. YG developed the extractive search concept and the SPIKE engine and oversees the technical parts of the project. HT wrote the code and UI for the knowledgebase app with guidance from YS. YS supervised the project.

## Notes

https://spike-kbc.apps.allenai.org

### Competing Interest Statement

The authors have declared no competing interest.

