## Supplementary Information for "Rapid Knowledgebase Construction and Hypotheses Generation Using Extractive Literature Search"

**for**

Yoav Goldberg

#### Contact Information

Dr. Yosi Shamay

Cancer Nanomedicine and Nanoinformatics Lab,

Biomedical Engineering Faculty, Technion – Israel Institute of Technology, Haifa, Israel

### Supplementary Figures

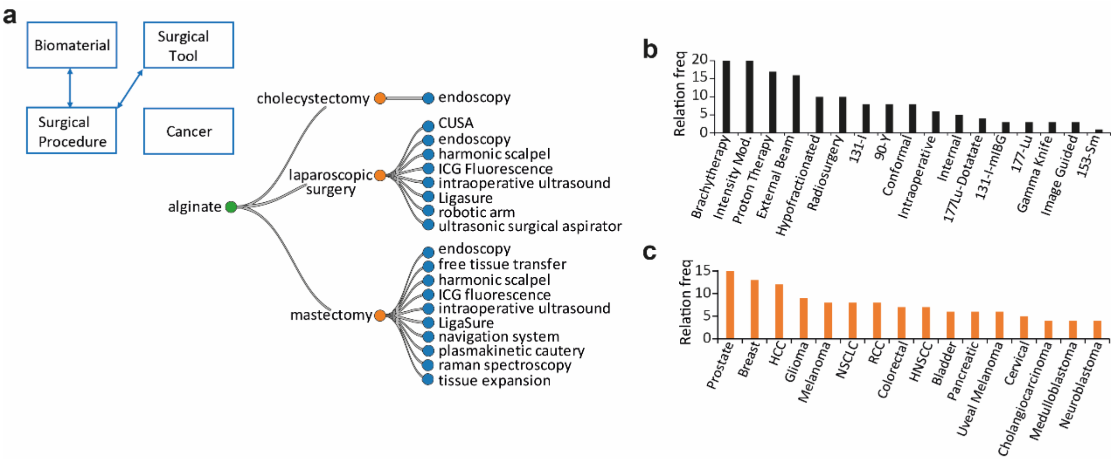


**Supplementary Fig 1. a)** example of a structured hypothesis starting from alginate and ending with surgical tools. **b)** analysis of number of relations between cancers and radiotherapies. **c)** analysis of number of relations between cancers and radiotherapies.

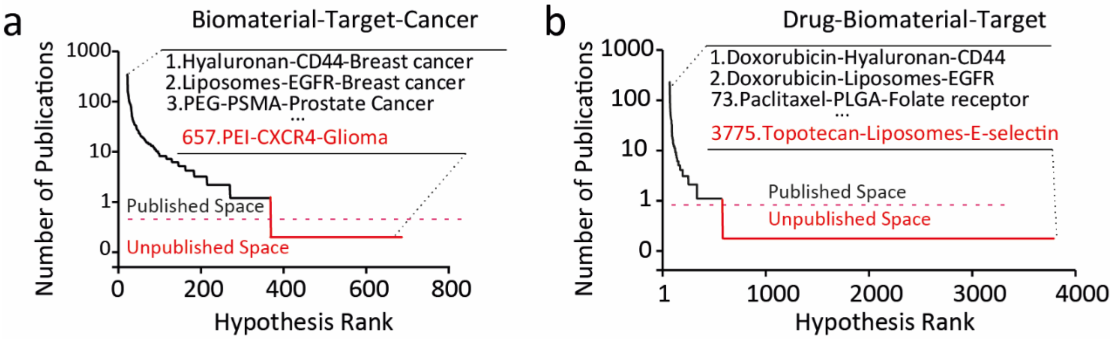


**Supplementary Fig 2. a)** Cancer-Target-Ligand based hypothesis combinations ranked by the number of publications in PubMed. **b)** Drug-Biomaterial-Target based hypothesis combinations ranked by the number of publications in PubMed

**Supplementary Table S1.** Biomedical entities mined from leading reviews in nanomedicine, used as keywords for the Bioportal' ontology recommender. words are in the order of appearance in text.

| **Manzari, M. T. et al. Targeted drug delivery strategies for precision medicines. Nat. Rev. Mater.** | **Mitchell, M. J.** *et al.* Engineering precision nanoparticles for drug delivery. *Nat. Rev. Drug Discov.* **20**, 101–124 (2021). | **Shi, J., Kantoff, P. W., Wooster, R. & Farokhzad, O. C. Cancer nanomedicine: progress, challenges and opportunities.** *Nat. Rev. Cancer* **17**, 20–37 (2017). |
|---|---|---|
| AKT | liposomes | Acute lymphoblastic leukemia |
| AMPK | lipid nanoparticle | acute myeloid leukemia |
| asialoglycoprotein receptor | LNP | adrenocortical carcinoma |
| AXL | lipid | albumin nanoparticles |
| Bcl-2 | phospho-lipids | alendronate |
| BCR- ABL | cholesterol | BIND-014 |
| BRAF | Peg | bortezomib |
| caveolae | nanocapsule | breast cancer |
| CD31/PECAM1 | nanosphere | Camptothecin |
| CDK | polymersome | Carbon |
| CDK4 | micelle | CD47 |
| CDK6 | dendrimer | cisplatin |
| CHK1 | amphiphilic block copolymer | adenocarcinoma |
| E- selectin | poly(ethylene glycol) | colorectal cancer |
| FAK | poly(dimethylsiloxane) | Combretastatin |
| FGFR1 | PDMS | CRGDK |
| FLT3 | poly(ethylenimine) | curcumin |
| HER2 | PEI | cytarabine |
| MAPK | poly(amidoamine) | daunorubicin |
| MEK | PAMAM | dendrimers |
| MER | Polyelectrolyte | B-cells |
| P- selectin | gold | bisphosphonates |
| PDGFR | iron | dextran |
| PECAM1 | silica | diffuse large B-cell lymphoma |
| PI3K | AuNP | cancer cells |
| PI3Kα | Iron oxide | disulfiram |
| PSMA | magnetite | Docetaxel |
| RAF | Magnetic iron oxide | doxorubicin |
| RTK | Fe3 | EC |
| TAM | O4 | EGFR |
| TYRO3 | maghemite | endothelial cells |
| VEGFR | Fe2 O3 | ERBB2 |
| cancer cells | calcium phosphate | erythrocytes |
| endothelial cells | mesoporous silica | Exosomes |
| stromal cells | Quantum dots | ferumoxytol |
| fibroblasts | silicon | fibroblasts |
| T-cells | PLGA | floxuridine |
| bisphosphonates | PBAE | folate receptor |
| RGD | PVC | galactose |

| **Manzari, M. T. et al. Targeted drug delivery strategies for precision medicines. Nat. Rev. Mater.** | **Mitchell, M. J.** *et al.* Engineering precision nanoparticles for drug delivery. *Nat. Rev. Drug Discov.* **20**, 101–124 (2021). | **Shi, J., Kantoff, P. W., Wooster, R. & Farokhzad, O. C. Cancer nanomedicine: progress, challenges and opportunities.** *Nat. Rev. Cancer* **17**, 20–37 (2017). |
|---|---|---|
| nanocrystal | poly(acrylamide- co- methacrylic acid) | Gastric cancer |
| Liposome | nanogel | glioblastoma |
| PLA | lipoplexes | Glioma |
| nanoparticles | thiolated hyaluronic acid | gold nanoshell |
| Idelalisib | N- acetylcysteine | graphene |
| Sorafenib | Polymer micelle Nanosphere | hafnium oxide |
| Abemaciclib | spherical NP | Head and neck cancer |
| gefitinib | pH- responsive NP | hepatocellular carcinoma |
| Cabozantinib | poly(β- amino- ester) | hepatocytes |
| alpelisib | lipofectamine | HER2 |
| Copanlisib | polyethylene imine | hydrazinocurcumin |
| Erlotinib | jetPEI | IL-2 |
| Trametinib | PAA | integrin |
| Dabrafenib | poly(amido amine) | irinotecan |
| Ceritinib | PLL | iron oxide |
| Lenvatinib | polylysine | Kaposi sarcoma |
| Buparlisib | cyclodextrins | leukocytes |
| SP600125 | poly(β- amino esters) | Lewis lung carcinoma |
| cisplatin | polyethylene oxide | Liposome |
| Midostaurin | hyaluronic acid | liver cancer |
| Bevacizumab | LNPs | lung cancer |
| Ponatinib | EGFR | macrophages |
| Nintedanib | epidermal growth factor receptor | magnetic NP |
| lapatinib | CAV1 | mannose |
| Sirolimus | clathrin | mannose receptor |
| Venetoclax | collagen | Melanoma |
| Bosutinib | transferrin receptor | Mesenchymal stem cell |
| Olaparib | endosome | mesoporous silicon |
| Axitinib | integrins | micelles |
| vandetanib | folate receptor | MUC1 |
| MEK163 | C- type lectin receptor | multiple myeloma |
| Afatinib | lectin receptors | myeloma |
| AZD2811 | DEC-205 | nano diamond |
| Crizotinib | CLEC9A | nanorod |
| doxorubicin | CD19 | nanoworm |
| GSK2256098 | scavenger receptor class B1 | Nanocrystal |
| Imatinib | SRB1 | nanoparticles |
| LY2606368 | PD1 | Neocarzinostatin |
| paclitaxel | CD3 | non-Hodgkin lymphoma |
| Pazopanib | THY1 | non-small-cell lung cancer |
| PD0325901 | CD90 | oesophageal adenocarcinoma |
| rapamycin | PDL1 | oligonucleotide |

| **Manzari, M. T. et al. Targeted drug delivery strategies for precision medicines. Nat. Rev. Mater.** | **Mitchell, M. J.** *et al.* Engineering precision nanoparticles for drug delivery. *Nat. Rev. Drug Discov.* **20**, 101–124 (2021). | **Shi, J., Kantoff, P. W., Wooster, R. & Farokhzad, O. C. Cancer nanomedicine: progress, challenges and opportunities.** *Nat. Rev. Cancer* **17**, 20–37 (2017). |
|---|---|---|
| regorafenib | CTLA4 | Osteoblast |
| SB202190 | T-cell | osteosarcoma |
| sonidegib | cancer cell | ovarian cancer |
| Sunitinib | intestinal epithelial cells | Oxaliplatin |
| vemurafenib | non- phagocytic cells | paclitaxel |
| Vismodegib | fibroblast cells | pancreatic cancer |
| B-cells | epithelial cells | PEComa |
| Dendrimer | amniotic stem cells | PEG |
| EGFR | macrophage | peritoneal cancer |
| folate receptor | leukocyte | peritoneal immune cells |
| galactose | erythrocyte | platinum |
| transferrin receptor | dendritic cells | pleural mesothelioma |
| macrophages | macrophages | PLA |
| PEG | dendritic cells as | PLGA-PEG |
| Silica | B-cells | PEI |
| | APCs | Polymeric micelles |
| | CREKA | polystyrene |
| | Mannose | prostate cancer |
| | galactose | Protein NPs |
| | dextran | PSMA |
| | sialoadhesin | pyrolipid |
| | Lyp1 | quantum dots |
| | anti-PD1 | Rapamycin |
| | anti-CD2 | renal cancer |
| | anti-CD4 | renal cell carcinoma |
| | sialic acid- binding immunoglobulin- like lectin | RGD |
| | | RGPD |
| | | sarcoma |
| | | Silica |
| | | silver |
| | | small cell lung cancer |
| | | soft tissue sarcoma |
| | | stromal cells |
| | | T-cells |
| | | Thermosensitive liposomes |
| | | thrombocytes |
| | | TNF |
| | | TRAIL |
| | | transferrin receptor |
| | | Tumour cell |
| | | VEGFR1 |

| **Manzari, M. T. et al. Targeted drug delivery strategies for precision medicines. Nat. Rev. Mater.** | **Mitchell, M. J.** *et al.* Engineering precision nanoparticles for drug delivery. *Nat. Rev. Drug Discov.* **20**, 101–124 (2021). | **Shi, J., Kantoff, P. W., Wooster, R. & Farokhzad, O. C. Cancer nanomedicine: progress, challenges and opportunities.** *Nat. Rev. Cancer* **17**, 20–37 (2017). |
|---|---|---|
| | | vincristine |
| | | Viral NPs |
| | | Vorinostat |
| | | yittrium-90 |

Link to Supplementary Video S1:

https://www.youtube.com/watch?v=zRxnNfdQNF0&t=791s
